# GAP-MS: Automated validation of gene predictions using integrated mass spectrometry evidence

**DOI:** 10.64898/2026.03.17.712294

**Authors:** Qussai Abbas, Mathias Wilhelm, Bernhard Kuster, Dimitri Frishman

## Abstract

Accurate genome annotation is fundamental to modern biology, yet distinguishing authentic protein-coding sequences from prediction artifacts remains challenging, particularly in complex plant genomes where automated methods are error-prone and manual curation is rarely feasible due to prohibitive time and costs. Here, we present GAP-MS (**G**ene model **A**ssessment using **P**eptides from **M**ass **S**pectrometry), an automated proteogenomic pipeline that leverages mass spectrometry evidence to validate the protein-level accuracy of predicted gene models. Applied across 9 major crop species, GAP-MS consistently improved the prediction precision for four widely used gene prediction tools. In addition to filtering likely erroneous models, the pipeline identified hundreds of candidate protein-coding loci absent from current standard reference annotations. These peptide-supported loci were further verified by transcriptional evidence, well-supported functional annotations, and high coding-potential scores. Together, these results demonstrate that direct proteomic evidence can help resolve annotation ambiguities, define high-confidence peptide-supported reference proteomes, and uncover overlooked protein-coding genes, while facilitating the identification of sequences that may require further investigation.

## 2. Introduction

High-throughput sequencing technologies and advances in assembly algorithms have led to a rapid increase in the number and quality of genome assemblies across the tree of life. Large-scale initiatives such as the Earth BioGenome Project [1] and affiliated efforts, including Darwin Tree of Life [2], the 10,000 Plant Genome Sequencing Project (10KP) [3], and the European Reference Genome Atlas [4], aim to generate reference genomes for all known eukaryotic species.

In contrast, genome annotation has progressed more slowly, and accurate prediction of protein-coding genes, their structures and isoforms remains a major bottleneck in turning raw assemblies into biologically interpretable resources [5, 6]. Computational gene prediction has therefore become a central focus, with a growing ecosystem of tools that combine *ab initio* models, comparative genomics and transcriptomic evidence to infer gene models. Early gene prediction tools for eukaryotes, including GeneMark [7], Genscan [8], and later Augustus [9], relied primarily on intrinsic sequence signals such as coding potential, splice-site motifs, and other statistical properties of genomic DNA to identify candidate coding regions. As annotation strategies evolved, pipelines began incorporating external evidence sources, including protein homology (e.g., orthologous genes) and experimental transcript data. Initially derived from expressed sequence tags (ESTs) and later from RNA-seq experiments, these data substantially improved gene model accuracy. Modern annotation frameworks such as Maker [10], Braker1/2/3 [11-13], the Gnomon system [14], and the Ensembl gene annotation pipeline [15] combine these heterogeneous evidence types to generate more reliable annotations. More recently, deep learning–based approaches, including Helixer [16] and Annevo [17], have emerged, applying neural network models to directly learn complex patterns in genomic sequences and further improve gene structure prediction.

Despite substantial progress, current pipelines still suffer from false positives and false negatives, fragmented gene models and incompletely resolved alternative splice variants, particularly in complex eukaryotic genomes [18] and crop species with large, repetitive, or polyploid genomes [19]. As a result, many predicted genes lack direct experimental support at the protein level, and a significant fraction of the proteome remains unannotated or misannotated [20]. These inaccuracies accumulate rapidly in public databases, propagating errors across subsequent analyses and tools that rely on these resources, which underscores the need for orthogonal validation strategies to filter and refine predicted gene models systematically.

At the same time, mass spectrometry–based proteomics is undergoing a parallel expansion in both scale and coverage, driven by large coordinated data-generation initiatives. One such effort is the crop proteome atlas within the “Proteomes that Feed the World” project [21], which aims to chart the proteomes of the 100 most important crops for human nutrition, alongside the continued accumulation of public datasets coordinated through the ProteomeXchange consortium [22].

Proteomics data offers a powerful and complementary line of evidence to address genome annotation errors. By directly detecting peptides derived from translated proteins, proteomics can provide physical confirmation of protein expression, validate predicted coding sequences and refine gene structures, including exon–intron boundaries and alternative start/stop sites [23]. Proteogenomic approaches have already demonstrated their value in prokaryotes, where relatively compact genomes facilitate comprehensive searches and genome-wide validation of gene models [24]. Early studies also showed that this approach can be applied to large, complex eukaryotic genomes, enabling direct protein identification from raw genomic sequence data [25]. Nevertheless, in eukaryotes, its applications have largely been restricted to individual species or focused studies [26, 27], and systematic large-scale integration of proteomics into routine genome annotation remains uncommon.

This imbalance highlights a critical methodological gap. Genome sequencing and mass spectrometry are now both high throughput and widely accessible, but the field still lacks robust, scalable and automated pipelines to exploit proteomics data for genome annotation at scale. Developing such pipelines, capable of systematically assessing predicted gene models in an automated and quality-controlled manner, will be essential to fully realize the potential of proteomics as an orthogonal source of evidence for eukaryotic genome annotation.

Here, we introduce GAP-MS, an automated proteogenomic pipeline designed to evaluate the protein-level support of predicted gene models by integrating mass spectrometry–derived peptide evidence. We apply GAP-MS across 9 major crop species, enabling large-scale, comparative assessment of genome annotations and providing direct experimental support for translated gene products.

## 3. Materials and Methods

### 3.1 Data retrieval

#### 3.1.1 Plant genomic data selection

Genome assemblies and annotation data for the top 100 crops essential for human nutrition were compiled from GenBank [28], following the procedures outlined in our previous work [19]. Briefly, the list of major food crops was obtained from the Food and Agriculture Organization (FAO) of the United Nations [29], ranked by annual production, and for each crop the most recent genome assembly and associated metadata were retrieved.

From this initial set of 100 crops, species were retained only if a RefSeq assembly and the corresponding GFF reference annotation were available and the assembly status was complete or chromosome level. RefSeq annotations were used as standardized reference sets for comparative benchmarking, providing a consistent baseline across species. Because reference annotations may still contain missing, fragmented, or incorrectly merged gene models, agreement with RefSeq was interpreted as concordance with the current community reference rather than absolute biological accuracy. To further ensure assembly and annotation quality, we retained only genomes generated with long-read sequencing (PacBio or Oxford Nanopore) and released after 2020. Gap content was required to be ≤1%, calculated as ((total sequence length – total ungapped length) / total sequence length × 100%), and only species annotated with NCBI Eukaryotic Genome Annotation Pipeline version ≥9.0 were included. Assembly and gene-model completeness were evaluated using BUSCO v5.2.2 [30] with the embryophyta_odb10 lineage dataset, and both the assembly and its annotation were required to achieve ≥98% complete (single-copy + duplicated) BUSCOs. We also assessed intergenic and repetitive-region contiguity using the LTR Assembly Index (LAI) [31], applying a threshold ≥10. Genomes failing any of these criteria were excluded.

This filtering strategy retained 18 crops with high-quality assemblies and annotations (Supplementary Table 1), from which we selected 9 crops with available mass spectrometry data at the time of the study. These 9 species include both monocots (e.g., maize and barley) and dicots (e.g., tomato and apple), span substantial phylogenetic diversity across the plant kingdom (Supplementary Fig. 1), and served as our benchmark datasets for evaluating gene prediction tools.

#### 3.1.2 Gene prediction

Gene prediction was conducted using four modern computational tools. Braker2 [12], Galba [30], and Helixer [16] were selected based on their superior accuracy in our previous benchmarking study [19]. Additionally, we included Annevo [17], a recently developed *ab initio* gene prediction tool, due to its reported high prediction accuracy. Braker2 and Galba utilize diverse protein databases to align proteins to the genome, thereby generating essential hints for Augustus [9] to predict gene models and infer multiple isoforms per gene. In contrast, Helixer employs a deep neural network with a bidirectional long short-term memory (LSTM) architecture to predict genes, typically yielding a single isoform per gene. Annevo applies a genomic language model that captures long-range sequence dependencies and joint evolutionary relationships across diverse genomes. All tools were executed using their default parameter settings.

#### 3.1.3 RNA-seq data collection

RNA-seq datasets for the 9 crops were retrieved from the Sequence Read Archive [31] using VARUS [32]. Reads were aligned to the corresponding genome assemblies with HISAT2 [33]. Aligned reads were converted to BAM format, sorted by genomic coordinates, and indexed for downstream analyses.

#### 3.1.4 Proteomics sample preparation and LC–MS/MS analysis

Plant tissues from the nine crop species were processed using the workflow described previously [34]. Briefly, proteins were extracted from multiple tissues by TCA/acetone precipitation followed by SDS-based solubilization and phenol extraction to remove metabolites and contaminants. The protein pellets were further purified with ammonium acetate in methanol, washed, re-solubilized, and quantified using a BCA assay. For proteomic analysis, 200 μg of protein per sample was subjected to automated SP3 bead-based cleanup and digestion using Sera-Mag magnetic beads, including reduction, alkylation, and overnight tryptic digestion, followed by peptide desalting. To increase proteome coverage, peptides were fractionated by basic pH reversed-phase chromatography into multiple fractions. Each fraction was then analyzed by micro-flow LC-MS/MS on an Orbitrap Eclipse Tribrid mass spectrometer in data-dependent acquisition mode, enabling high-resolution peptide identification and quantification. The mass spectrometry proteomics data have been deposited to the ProteomeXchange Consortium via the PRIDE [35] partner repository with the dataset identifier PXD076526.

#### 3.1.5 Peptide and protein identification

Raw label-free MS/MS datasets were analyzed using the FragPipe v23.1 pipeline [36]. Protein FASTA databases corresponding to (i) RefSeq reference annotations and (ii) predicted proteomes from Braker2, Galba, Helixer, and Annevo were prepared as independent search spaces using Philosopher v5.1.2 [37], including reversed-sequence decoys and common LC–MS/MS contaminants from the cRAP database [38]. Spectra were searched with MSFragger v4.3 [39] using trypsin as protease and allowing up to two missed cleavages. Peptide–spectrum matches (PSM) were rescored with MSBooster v1.3.17 [40], leveraging Koina [41] to obtain Prosit deep-learning predictions for fragment ion intensity (Prosit_2020_intensity_HCD) and retention time (Prosit_2019_irt) [42]. PSM validation was performed with Percolator v3.7.1 at a 1% False Discovery Rate (FDR) threshold [43], followed by protein inference and FDR filtering using Philosopher with a 1% protein-level FDR.

#### 3.1.6 Evaluation of peptide identification

To evaluate how well predicted proteomes derived by different gene annotation tools support peptide identification, each predicted proteome from Braker2, Galba, Helixer and Annevo was tested independently as a search space for peptide identification. The resulting peptide sets were then compared to those obtained using the reference RefSeq annotation. For each species and annotation, we quantified (i) the total number of identified peptides, (ii) the number and fraction of peptides shared with RefSeq, and (iii) the number of peptides uniquely identified using the predicted proteins (Fig. 1).

**Figure 1.**
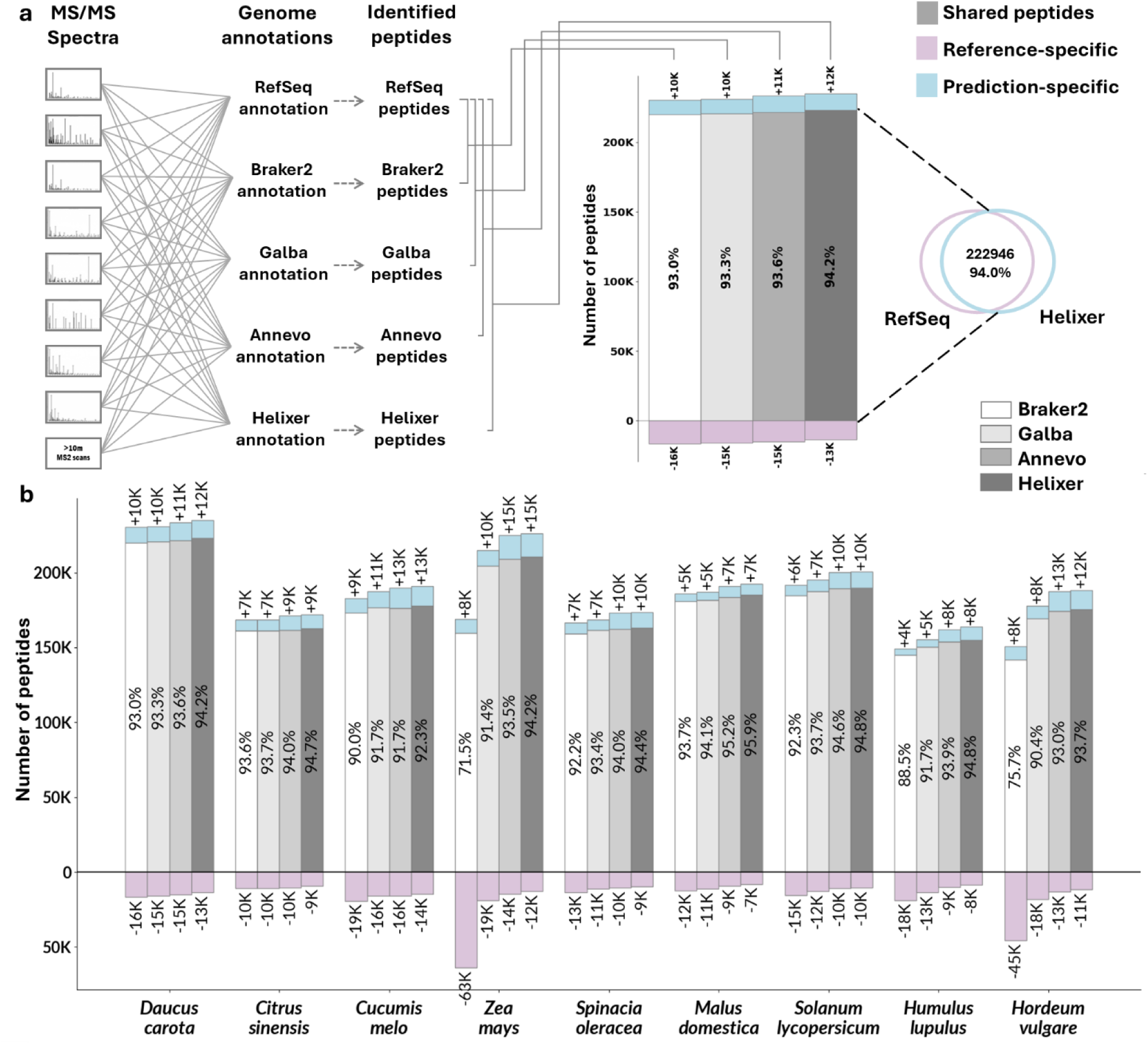
Comparative peptide identification rates of gene prediction tools across 9 crop species. **(a)** Schematic of the evaluation workflow: Raw MS/MS spectra are searched in parallel against the RefSeq annotation and predicted gene models (Braker2, Galba, Annevo, Helixer). The resulting identified peptides are compared to categorize results: <u>Shared peptides</u> (gray; consensus), <u>Reference-speci</u>fi<u>c peptides</u> (pink; missed by prediction), and <u>Prediction-specific</u> <u>peptides</u> (blue; putative novel genes). The Venn diagram illustrates this overlap for a representative comparison. **(b)** Stacked bars represent the total peptide yield for each prediction tool across 9 species. Gray bars indicate agreement with the reference; pink bars (negative y-axis) indicate reference-specific peptides; and blue bars indicate peptides unique to the prediction. Numeric labels denote peptide counts (in thousands, K), and percentages within the bars denote the recovery rate, calculated as (𝑁_𝑆ℎ𝑎𝑟𝑒𝑑_/ 𝑁_𝑅𝑒𝑓𝑒𝑟𝑒𝑛𝑐𝑒_) × 100%, where 𝑁_𝑅𝑒𝑓𝑒𝑟𝑒𝑛𝑐𝑒_ is the sum of shared and reference-specific peptides. The four bars for each species correspond to Braker2, Galba, Annevo, and Helixer, respectively (left to right).

### 3.2 GAP-MS pipeline

We developed GAP-MS, a pipeline that integrates mass-spectrometry–based proteomic evidence with in-silico predictions to evaluate and filter gene models. The pipeline is designed to work across different gene prediction tools and takes as input predicted gene models in GTF/GFF format, the corresponding genome assembly in FASTA format, and a list of mass-spectrometry–identified peptides in TXT format. As an optional input, RNA-seq alignment files in BAM format can also be provided. Using these inputs, GAP-MS extracts predicted protein sequences from gene models using gffread [44] and assigns confidence levels based on peptide-mapping evidence, thereby distinguishing peptide-supported coding sequences from probable annotation artifacts. When RNA-seq data are included, the pipeline uses StringTie3 [45] to assemble transcripts, TransDecoder to predict coding regions, and gffread [44] to translate them into predicted ORF proteins. These ORFs are then used to validate peptide-supported gene models at the transcriptional level, providing an additional layer of support. GAP-MS further enables comparative analyses against reference annotations, identification of novel protein-coding candidates, and detection of potentially misannotated gene models.

#### 3.2.1 Peptide mapping and feature engineering

GAP-MS begins by mapping the identified peptides from the mass spectrometry experiments to the protein sequences of the predicted gene models using Proteomapper v1.5 [46]. For each protein, a feature set is compiled that integrates sequence attributes with proteomic evidence. Sequence features include protein length, the number of splice sites, and the number of annotated isoforms. Proteomic features include the number of mapped peptides, sequence coverage (the percentage of residues overlapped by identified peptides, typically 10 to 40% and occasionally above 70% for highly expressed proteins), peptide uniqueness at the protein and gene levels, and the position of each peptide within the protein. Peptides are classified as gene-specific (mapping to any isoform of the same gene), protein-specific (mapping uniquely to a single protein isoform), internal (entirely within one exon), splice (spanning an exon–exon junction), N-terminal (mapping to the start of the protein), and C-terminal (mapping to the region immediately upstream of the stop codon). The resulting feature matrix is used to train the classification model described below.

#### 3.2.2 Classification of proteins into confidence groups

Based on the extracted features, gene models were partitioned into three sets: high-confidence, low-confidence, and unlabeled. High-confidence proteins were defined by robust proteomic support, meeting at least one of the following stringent criteria: the presence of two or more unique protein-specific peptides, sequence coverage of at least 80%, detection of both N-terminal and C-terminal peptides, or complete support for all annotated splice sites by corresponding splice peptides. In contrast, low-confidence proteins were characterized by limited peptide evidence, specifically exhibiting fewer than two mapped peptides and an absence of gene-specific peptides. Proteins with intermediate evidence were designated as unlabeled.

To resolve the status of gene models with intermediate unlabeled evidence, a multivariate XGBoost classifier [47] was employed, capturing non-linear relationships between protein length, sequence coverage, peptide uniqueness, and peptide location that rule-based filtering would systematically misclassify. Accordingly, the high-confidence and low-confidence sets served as positive and negative training examples, respectively. The resulting model was applied to the unlabeled set to assign a final status of verified or dismissed. The classifier was optimized using randomized hyperparameter search with stratified 5-fold cross-validation and ROC-AUC scoring. Details on model training, performance evaluation, and score cutoffs are provided in Supplementary File 1.

#### 3.2.3 Model interpretation and validation

To elucidate the contributions of individual features to protein classification, SHAP (SHapley Additive exPlanations) analysis was employed [48]. This analysis revealed the most impactful features influencing model predictions, thereby validating the biological relevance of our chosen metrics.

### 3.3 Assessment of GAP-MS impact on gene prediction accuracy

The accuracy of Braker2, Galba, Helixer, and Annevo (both with and without applying GAP-MS) was evaluated at the exon, intron chain, and transcript levels by comparing their boundaries to those in the RefSeq reference annotations of 9 crops using Gffcompare v0.12.10 [44]. Specifically, exon level accuracy required exact matching of start and end coordinates for individual coding exons. Intron chain level accuracy evaluated the precise sequence of internal splice junctions within a transcript, disregarding the outer transcription start and end sites. Transcript level accuracy imposed the strictest criteria, requiring an identical match for all exon boundaries throughout the entire gene model [44]. Accuracy metrics included recall (the ability to detect true positive genes), precision (the ability to eliminate false positives), and F1 score (the harmonic mean of recall and precision).

Due to the non-normal distribution of the data, as confirmed by the Shapiro-Wilk test, non-parametric statistical methods were employed. Specifically, the Kruskal-Wallis test was used to compare the distribution of each metric across the tools. For pairwise comparisons, the Mann-Whitney U test was applied to identify significant differences between individual tools.

### 3.4 Identification of novel gene loci

Gene models predicted by gene prediction tools and supported by peptide evidence in GAP-MS, but absent from the RefSeq reference annotations, were subjected to additional validation and characterization. To identify high-confidence novel coding regions, we cross-referenced the GAP-MS supported gene models against the RefSeq models using gffcompare. Transcripts assigned class code ‘u’ (intergenic) or ‘x’ (antisense) were designated as novel gene loci, while those with class code ‘j’ were identified as novel isoforms of annotated genes.

#### 3.4.1 Transcriptional validation with RNA-seq

BAM files generated from RNA-seq alignments were used to provide transcriptional validation for peptide-supported gene models. Per-base coverage and exon–intron support were quantified using BEDTools [49]. For each novel peptide-supported gene model, coverage across the coding sequence was summarized. In parallel, Gffcompare was used to compare the overlap between the RNA-seq-derived ORF models and the peptide-supported gene models, enabling the selection of high-confidence novel candidate gene models supported by both proteomic and transcriptional evidence for further analysis.

#### 3.4.2 Assessment of protein coding potential

To further validate the coding potential of the novel gene models, the protein sequences were evaluated using Psauron [50]. This tool utilizes a machine learning model trained on over 1,000 plant and animal genomes to assign a probability score reflecting the likelihood that a query sequence represents a genuine protein-coding region. Predicted proteins were classified based on the tool’s binary output, where sequences labeled as true sequences were considered high-quality candidates distinguishable from spurious open reading frames.

#### 3.4.3 Functional annotation of protein sequences

Functional annotation of the novel peptide-supported proteins was performed with InterProScan v5 [51]. We recorded InterPro signatures, predicted domains, and associated Gene Ontology (GO) terms, and we flagged sequences lacking any domain assignment.

#### 3.4.4 Visualization of gene models

Gene models were visualized and inspected in the IGV genome browser [52]. Reference genome assemblies were loaded together with GFF annotation tracks for standard RefSeq gene models, raw gene prediction outputs, and the final GAP-MS–filtered gene sets. Transcriptional support was evaluated using HISAT2 BAM alignments to visualize read coverage depth and splice junctions. Proteomic evidence produced by GAP-MS (BED format) was mapped to genomic coordinates and displayed as custom tracks.

## 4. Results

### 4.1 Benchmarking of gene prediction tools using proteomics data

In this study, we evaluated how well the predicted proteins derived from gene prediction tools perform as search spaces for peptide identification in proteomics analyses. Parallel database searches were performed for nine crop species, matching raw MS/MS spectra against peptide sequences derived from these predicted protein sets. This assessment quantifies the concordance between peptides identified using predicted sequences from four gene prediction tools (Braker2, Galba, Annevo, and Helixer) versus those derived from standard RefSeq reference annotations. A high degree of overlap confirms that the prediction accurately captures the expressed proteome. Based on this comparison, the sets of identified peptides were categorized into three distinct groups: **shared peptides** (consensus support), **reference-specific peptides** (coding regions missed by the tool), and **prediction-specific peptides** (potential novel coding sequences absent from the RefSeq annotation) (Fig. 1a).

The predicted proteomes generally showed high concordance with RefSeq reference datasets (Fig. 1b); shared peptides typically accounted for >90% of the total identified spectra. Helixer consistently demonstrated superior predictive accuracy, achieving the highest peptide recovery rates and the lowest number of missed reference peptides across all species. While “prediction-specific” peptides (potential novel coding sequences) ranged between 5,000 and 15,000 peptides per species, “reference-specific” peptides (missed by prediction) were generally more numerous, typically ranging from 8,000 to 19,000. Species-specific variations were also observed. Braker2 predictions for *Zea mays*, and *Hordeum vulgare* diverged significantly from the reference, exhibiting a marked loss of reference-specific peptides (ranging from 45,000 to 63,000), indicating reduced Braker2 predictive quality for these species.

Although prediction-specific peptides may partly result from differences in search-space size, database redundancy, altered peptide assignment under different database contexts, or false-positive prediction models, their relatively high numbers and strong agreement across prediction tools suggest that many represent genuine novel candidate genes absent from current reference annotations. By examining the overlap among prediction-specific peptides identified by different tools per species, we found that, on average, >70% were supported by at least two tools, 34% were shared across all four tools, and fewer than 30% were unique to a single tool (Supplementary Fig. 2). This pattern suggests systemic incompleteness in current RefSeq annotations rather than an artifact of the prediction tools. These findings highlight the utility of mass spectrometry data as an orthogonal metric for both benchmarking prediction tools and identifying gaps in existing reference annotations.

### 4.2 Development of a proteomic-guided classification model (GAP-MS)

To distinguish authentic protein-coding regions from annotation artifacts, we developed GAP-MS, a pipeline that integrates genomic features with direct proteomic evidence (Fig. 2a). For each predicted gene model, GAP-MS extracts a feature set that combines sequence attributes with quantitative mass spectrometry metrics. Species-specific high-confidence training sets were defined from gene models with strong peptide support, whereas low-confidence sets comprised models with limited peptide evidence. An XGBoost classifier was then trained independently for each species using the high- and low-confidence sets as positive and negative examples and applied to ambiguous proteins with intermediate proteomic evidence to assign a binary status of verified or dismissed (see *Methods*). The final set of peptide-supported proteins therefore includes both high-confidence models and predictions verified by the classifier. This classification is illustrated for *Daucus carota* (Fig. 2b), which highlights marked differences in reliability among the four prediction tools. Although all tools identified a similar core of 15,000 to 16,000 high-confidence proteins, their total outputs differed substantially. Braker2 and Galba generated the largest initial protein sets but also produced the highest numbers of non-validated models, with 31,743 and 24,198 proteins assigned to the low-confidence category, respectively. In contrast, Annevo and Helixer yielded fewer dismissed models and therefore higher precision. Annevo was the most conservative, producing only 6,931 low-confidence proteins.

**Figure 2.**
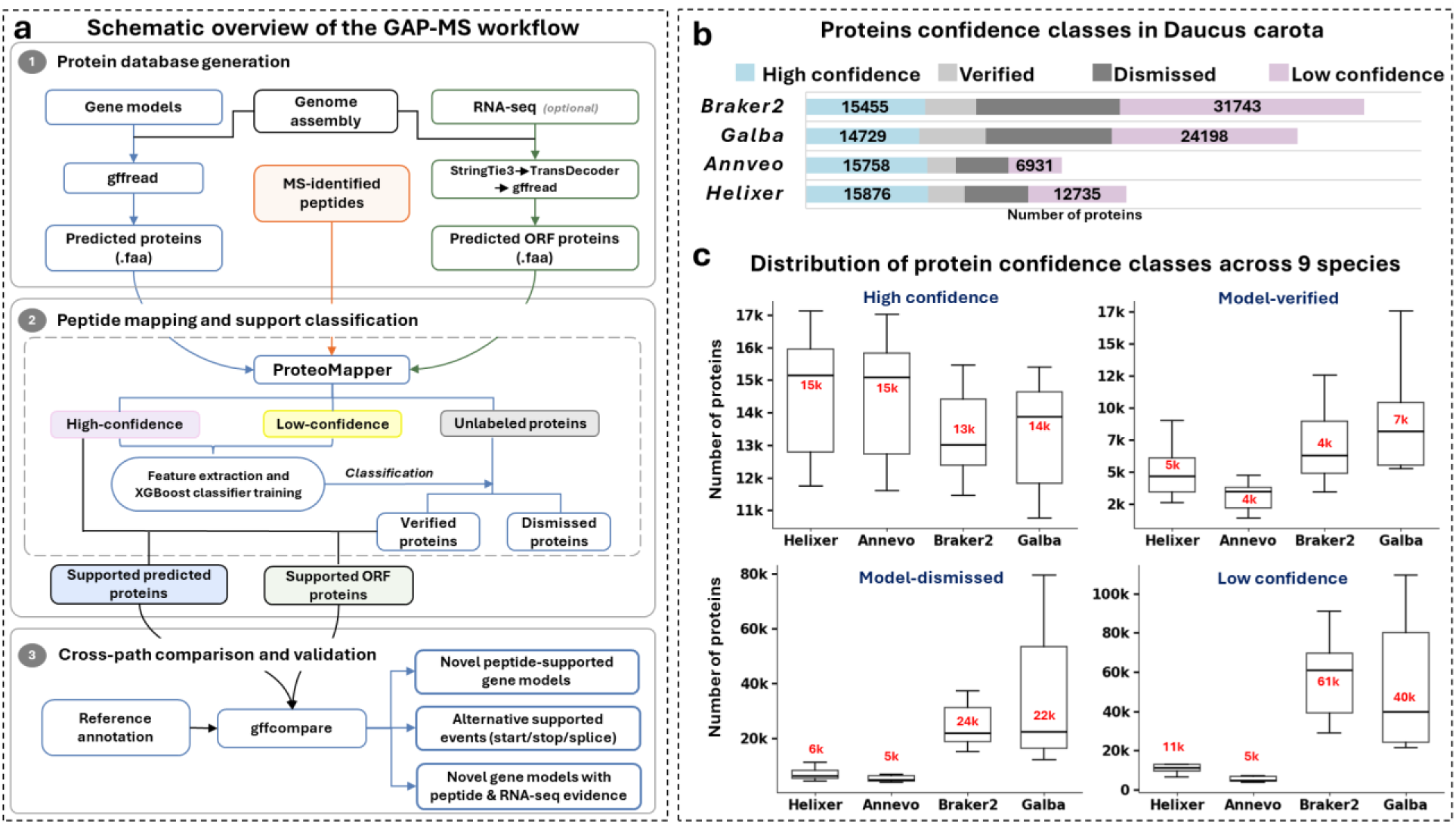
Design, training, and validation of the GAP-MS pipeline. **(a)** Schematic overview of the GAP-MS pipeline. Gene predictions are integrated with MS/MS peptide data to define training sets (high and low confidence) and train a species-specific XGBoost classifier for determining the support status of unlabeled proteins. (b) Classification profile for *Daucus carota*. Stacked bars display the absolute number of proteins assigned to high-confidence, verified, dismissed, and low-confidence categories for each prediction tool. (c) Distribution of protein confidence classes across 9 species. Box plots visualize the variation in protein counts for each classification category across the evaluated prediction tools. Center lines represent medians.

Expanding this analysis to the full dataset of 9 crops, proteomic support rates varied substantially across the prediction tools. The support percentage was calculated by dividing the total number of peptide-supported proteins by the total number of predicted proteins. This evaluation revealed that deep learning-based tools maintained relatively balanced distributions, with 63% and 51% of the total predictions classified as supported for Annevo and Helixer, respectively. Conversely, Braker2 and Galba exhibited heavily skewed distributions, with only 14% and 20% of their gene models receiving proteomic support (Supplementary Fig. 3).

This difference is primarily driven by the total number of predicted proteins. Braker2 and Galba produced substantially larger predicted protein sets (averaging 150k and 100k per species, respectively) compared to the more conservative outputs of Annevo and Helixer (averaging 29k and 36k per species, respectively). However, despite the lower support percentages, the absolute number of supported proteins yielded by Braker2 and Galba (average ∼20,000) was comparable to that of Annevo and Helixer (average ∼19,000–20,000). These patterns indicate that while Helixer and Annevo prioritize precision, homology-based approaches like Braker2 and Galba generate a substantial excess of unsupported potential artifacts to capture a similar core of authentic proteins (Fig. 2c).

Analysis of the peptide evidence underlying these classifications revealed that supported models were substantiated by high-quality features. On average, 15,094 proteins per crop were validated by protein-specific peptides across the four prediction tools, while 11,085 proteins received support from splice-spanning peptides. Complementary evidence was provided by terminal-specific peptides, with an average of 1,864 proteins supported by N-terminal peptides and 1,899 proteins exhibiting C-terminal peptide support per crop (Fig. 3a). Notably, Braker2 showed a clear deficiency in C-terminal peptide validation, with an average of only 447 supported proteins per crop, compared with approximately 2,300 in Galba, Annevo, and Helixer. This deficit at the 3′ end of gene models suggests that Braker2 has difficulty accurately defining stop codons or maintaining reading-frame consistency across terminal introns. Together, these results demonstrate that GAP-MS serves not only as a filtering approach but also as a practical diagnostic tool for identifying systematic structural errors in genome annotation pipelines.

**Figure 3.**
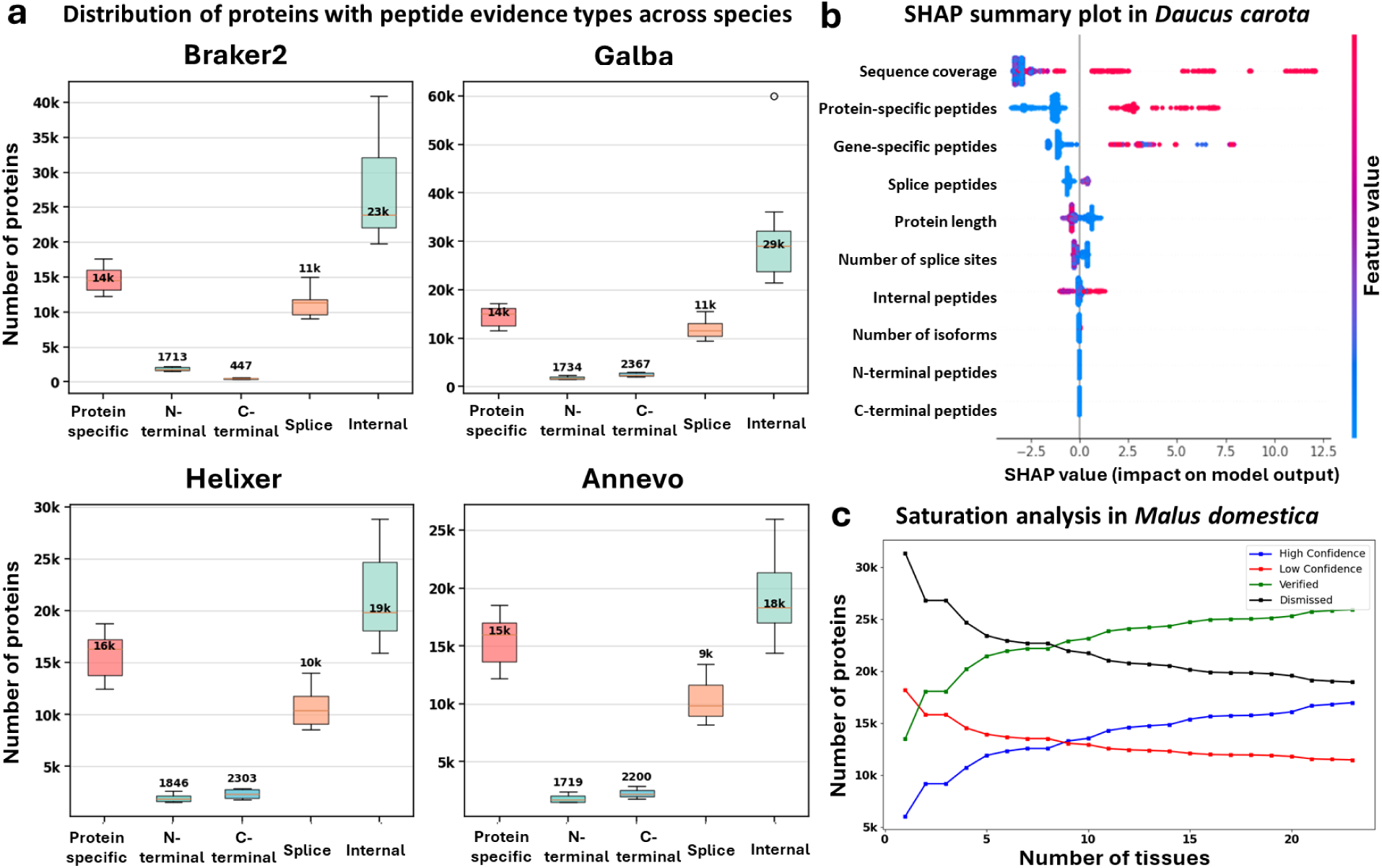
Characterization of peptide evidence and model interpretation. **(a)** Distribution of peptide evidence types across 9 crops. Box plots quantify the number of proteins supported by protein-specific, N-/C-terminal, splice-spanning, and internal peptides for each prediction tool. Center lines represent medians; box limits indicate the interquartile range. **(b)** SHAP feature importance analysis for Helixer predictions in *Daucus carota*. Features are ranked by their contribution to the model output. Each dot represents a protein, with color indicating the feature value (red = high, blue = low) and x-axis position denoting the impact on the classification probability (positive values favor "verified" status). **(c)** Proteomic saturation analysis in *Malus domestica*. The cumulative count of classified proteins is plotted against the number of merged tissue datasets. The trajectory shows a rapid initial accumulation of supported models followed by a plateau after approximately 15 tissues.

To understand which specific attributes most strongly influenced the distinction between verified and dismissed models, SHAP analysis was conducted, revealing that sequence coverage and the presence of protein-specific peptides consistently emerge as the primary drivers of the classification model across all species (one example in *Daucus carota* is shown in Fig. 3b).

Moreover, to evaluate the impact of proteomic depth on gene model validation by GAP-MS, a saturation analysis was performed on *Malus domestica,* selected for its comprehensive availability of 23 distinct tissue datasets. By cumulatively merging these tissue datasets (from 1 to 23), the count of peptide-supported gene models was tracked. The resulting curve shows a sharp initial increase between 1 and 5 tissues, where the inclusion of distinct datasets resulted in substantial gains in gene support. This confirms that multi-tissue atlases are essential for detecting hidden genes. Importantly, this trend decelerated after the 15th tissue, with the curve approaching a plateau that indicates proteomic saturation (Fig. 3c).

### 4.3 Improving precision through proteomic filtering

The performance of the four selected gene prediction tools (Braker2, Galba, Helixer, and Annevo) was evaluated across 9 crop species at the exon, intron chain, and transcript levels, both prior to and following filtering with GAP-MS. Across all tested tools, the application of GAP-MS consistently enhanced precision, with an expected tradeoff in recall (Fig. 4).

**Figure 4.**
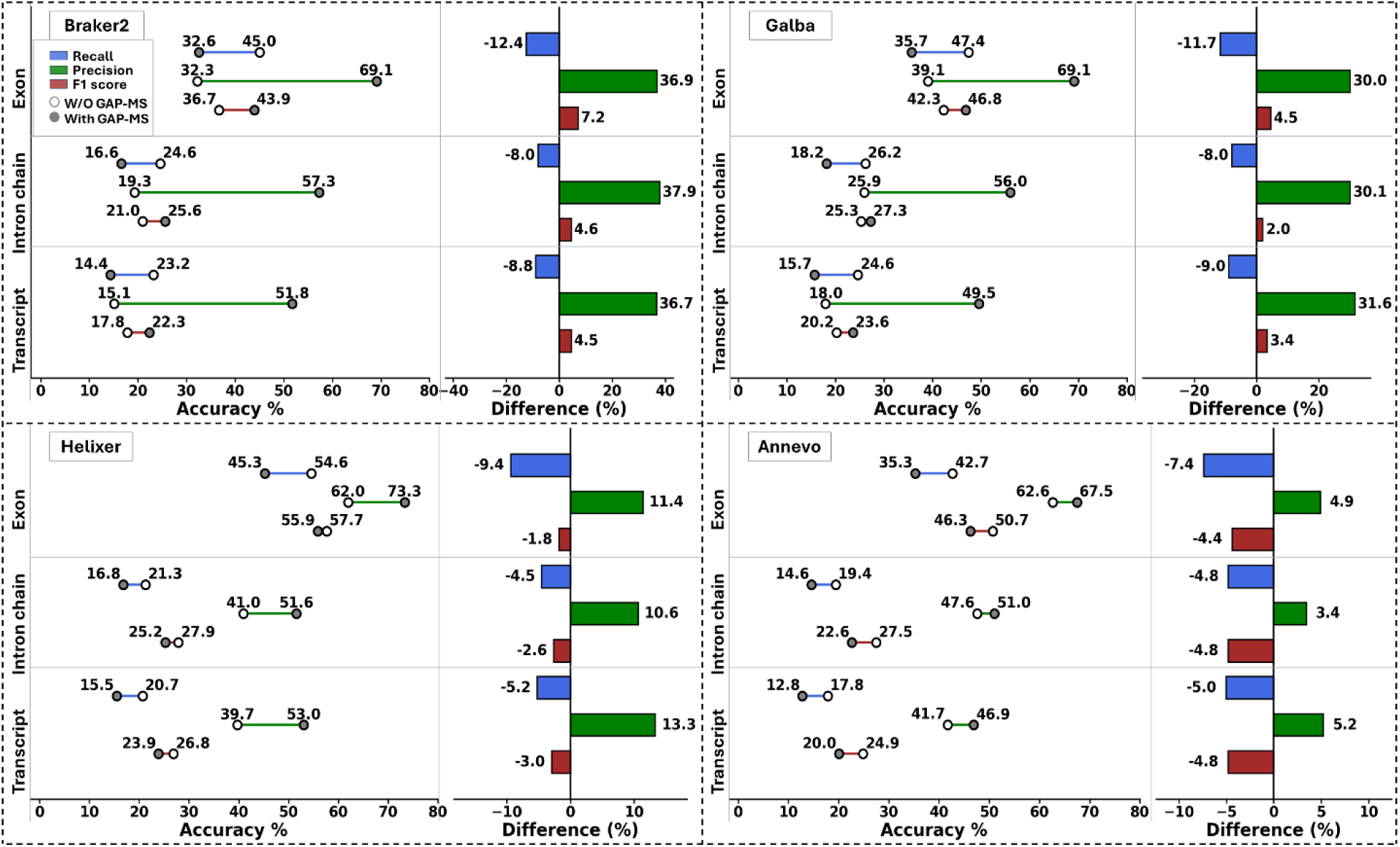
Comparative assessment of gene model accuracy before and after GAP-MS integration. Average accuracy metrics for Braker2, Galba, Helixer, and Annevo at three genomic levels (Exon, Intron chain, and Transcript) across 9 crop species. Left panels: Dumbbell plots illustrate the performance shift from the raw prediction (open white circles) to the GAP-MS refined set (filled grey circles). Colors denote specific metrics: Recall (blue), Precision (green), and F1 Score (red). Right panels: Bar charts quantify the performance differential, calculated as (GAP-MS metric – raw metric). Positive values (right-extending) indicate an increase in the metric, while negative values (left-extending) denote a reduction attributable to filtering.

The largest improvements were observed for Braker2 and Galba. These tools showed lower baseline precision (average initial transcript F1 scores of 18–20%) but a marked increase in precision after GAP-MS filtering, by approximately 37% for Braker2 and 31% for Galba across all feature levels. This increase came at the cost of sensitivity, with recall decreasing by approximately 8–12%. Nevertheless, the overall F1 score improved at all levels: transcript-level F1 scores increased by 4.5% for Braker2 and 3.4% for Galba, indicating that the removal of false positives outweighed the loss of true positives for these tools (Fig. 4). Deep-learning-based tools showed higher baseline accuracy, with initial average transcript F1 scores of approximately 25% for Annevo and 27% for Helixer. GAP-MS filtering further improved transcript-level precision by 13% in Helixer and 5% in Annevo. However, these gains were accompanied by reduced recall, resulting in modest decreases in transcript-level F1 scores of 3% for Helixer and 5% for Annevo.

Statistical analysis confirmed that the shifts in the accuracy scores post-GAP-MS were significant across all tools (Mann-Whitney U test, 𝑝 < 0.05). Collectively, these data demonstrate that GAP-MS serves as a rigorous specificity filter, most effectively enhancing the performance of tools prone to high false-positive rates.

### 4.4 Validation of novel peptide-supported genes

Reference genome annotations often miss genuine genes because of factors such as the absence of detectable homology, divergent codon usage, lineage-specific gene expansions, or masking of low-complexity regions [55, 56]. In addition, micro-exons of 30 bp or less are particularly difficult to detect because they frequently fall below statistical thresholds or are discarded during splice-graph construction [57]. Consequently, gene models predicted by annotation tools and supported by peptide evidence through GAP-MS represent strong candidates for authentic coding sequences missing from current reference annotations.

GAP-MS enables the recovery of these missing or misannotated coding regions by comparing peptide-supported predictions against user-defined reference annotations. This comparison identifies fully novel gene models, as well as refinements of existing annotations where proteomic evidence supports alternative translation start sites, alternative stop sites, or altered exon–intron structures. We therefore classified peptide-supported differences into four categories: novel genes, alternative splice structures, alternative start codons, and alternative stop codons, and summarized their distribution across tools and species (Fig. 5a).

**Figure 5.**
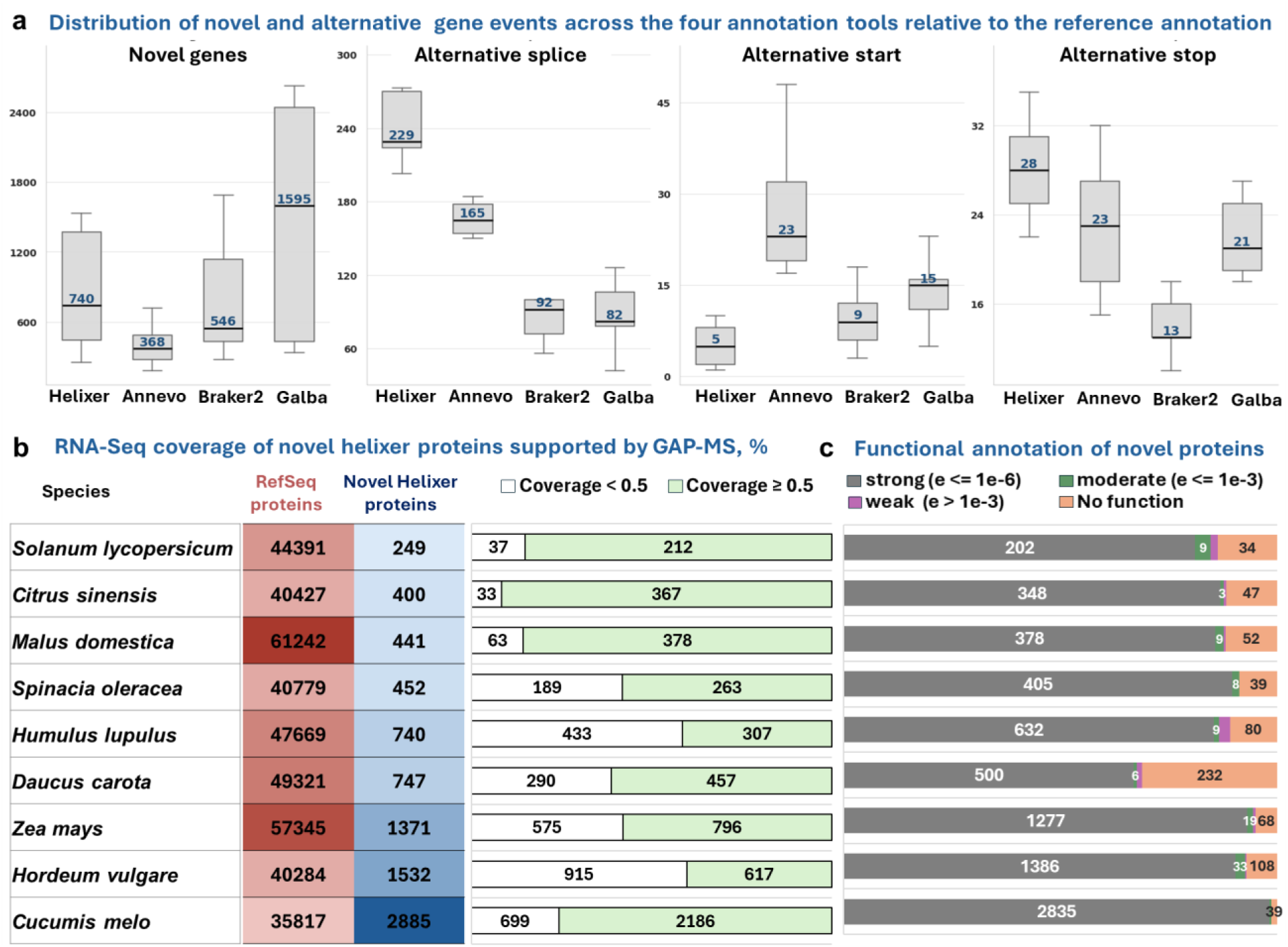
Transcriptional validation and functional characterization of novel peptide-supported gene models. **(a)** Distribution of peptide-supported annotation differences across the four gene prediction tools relative to the reference annotations, including novel genes, alternative splice events, alternative start sites, and alternative stop sites. **(b)** RNA-seq support analysis across 9 crop species. The heatmap compares the abundance of reference proteins (red) against novel peptide-supported Helixer predictions (blue). Horizontal bars quantify RNA-seq coding sequence coverage of the novel sets: high support (≥50% breadth, dark green) versus low support (<50% breadth, white). **(c)** Functional annotation profile. Stacked bars display the classification of novel proteins based on sequence homology to known domains, stratified by statistical significance (E-value): strong (≤ 10^−6^), moderate (≤ 10^−3^), weak (> 10^−3^), or no functional assignment.

Across the four annotation tools and nine crop species, peptide-supported novel genes represented the most abundant class of annotation difference relative to the reference annotations. Median novel gene counts were highest for Galba (1,595), followed by Helixer (740), Braker2 (546), and Annevo (368). In contrast, peptide-supported refinements of existing annotations occurred at substantially lower frequencies than novel gene discovery (Fig. 5a). Among structural refinements, alternative splice events were relatively more frequent, suggesting that exon–intron structure remains a major source of disagreement between reference annotations and peptide-supported predictions. By comparison, start- and stop-site corrections were less common, indicating that terminal boundary errors represent a smaller component of annotation incompleteness. Together, these results show that GAP-MS enables both novel gene discovery and targeted improvement of existing gene models. We next assessed the overlap of novel peptide-supported candidate genes across the four annotation tools. On average across species, approximately 60% of novel candidates were supported by at least two tools, including 13.5% detected by all four tools, whereas 40% were unique to a single tool (Supplementary Fig. 4).

Because Helixer showed the highest overall prediction accuracy and peptide recovery among the evaluated tools, we focused subsequent validation analyses on Helixer-derived novel proteins. Across the nine crop species, GAP-MS identified 8,817 novel peptide-supported proteins from Helixer predictions. The number of novel proteins varied substantially among species, ranging from 249 in Solanum lycopersicum to 2,885 in Cucumis melo (Fig. 5b). This variation was not explained by the size of the annotated RefSeq proteome, as the number of RefSeq protein-coding genes showed no significant correlation with the number of novel Helixer peptide-supported proteins (Pearson’s r = −0.36, p = 0.34; Supplementary Fig. 5).

To independently assess Helixer-predicted genes absent from RefSeq annotations, we examined RNA-seq read coverage breadth. As shown in Figure 5b, a substantial proportion of these proteins showed extensive transcriptional support (coverage ≥ 50%), and in most species high-coverage models outnumbered low-coverage ones. Thus, the RNA-seq analysis provides independent transcriptional support for these peptide-supported loci. Coverage density distributions further showed that, whereas low-confidence proteins peaked near zero coverage, novel peptide-supported proteins displayed a profile similar to that of the high-confidence set, with a strong skew toward full coverage (100%). This pattern is exemplified in *Solanum lycopersicum* (Supplementary Fig. 6), where 249 novel proteins showed high transcript coverage, compared with 37 showing low coverage. Cases in which low-confidence models showed high transcript coverage likely reflect the detection limits of mass spectrometry or condition-specific expression not captured in the sampled proteomes. Conversely, some novel proteins showed low RNA-seq coverage, which may help explain why these loci were previously overlooked in RefSeq annotation.

Furthermore, the coding potential of these novel sequences was independently verified using Psauron, a machine learning-based tool designed to distinguish genuine protein-coding regions from spurious open reading frames. Across the 9 crops, an average of 95% of the novel peptide-supported proteins were classified as true high-quality coding sequences (Supplementary Fig. 7), reinforcing that GAP-MS recovers authentic protein-coding sequences.

Finally, to assess the biological validity of the novel gene models recovered by GAP-MS, a systematic functional annotation of all novel Helixer-predicted, peptide-supported proteins across the 9 crop species was performed. Remarkably, over 95% of these novel sequences exhibited strong evidence of functional activity, characterized by significant homology to known protein domains or families (Fig. 5c). This high rate of functional conservation confirms that GAP-MS recovers evolutionarily conserved functional proteins.

### 4.5 Recovery of missing functional genes

To explore the biological significance of the recovered genes in greater depth, we focused on the *Malus domestica* proteome, where we identified over 7,000 peptides absent from the RefSeq annotation. Out of the 44,900 Helixer-predicted gene models, GAP-MS validated 25,030 peptide-supported models, which included 2,169 models with N-terminal peptide support and 2,778 with C-terminal support. Moreover, it recovered 441 novel peptide-supported gene models previously missing from the current RefSeq proteome.

The pipeline’s utility in resolving annotation gaps is exemplified by the recovery of a novel gene model functionally annotated as a C3HC4-type RING finger protein, a putative E3 ubiquitin ligase. While this specific gene is missing from the standard NCBI RefSeq annotation, GAP-MS identified a robust gene model supported by 88.3% RNA-seq breadth and three unique splice-spanning peptides (Fig. 6a). Crucially, this protein sequence overlaps with an existing entry in the Ensembl annotation, allowing GAP-MS to act as an independent proteomic validator that confirms the Ensembl model is correct and that the RefSeq exclusion represents a false negative.

**Figure 6.**
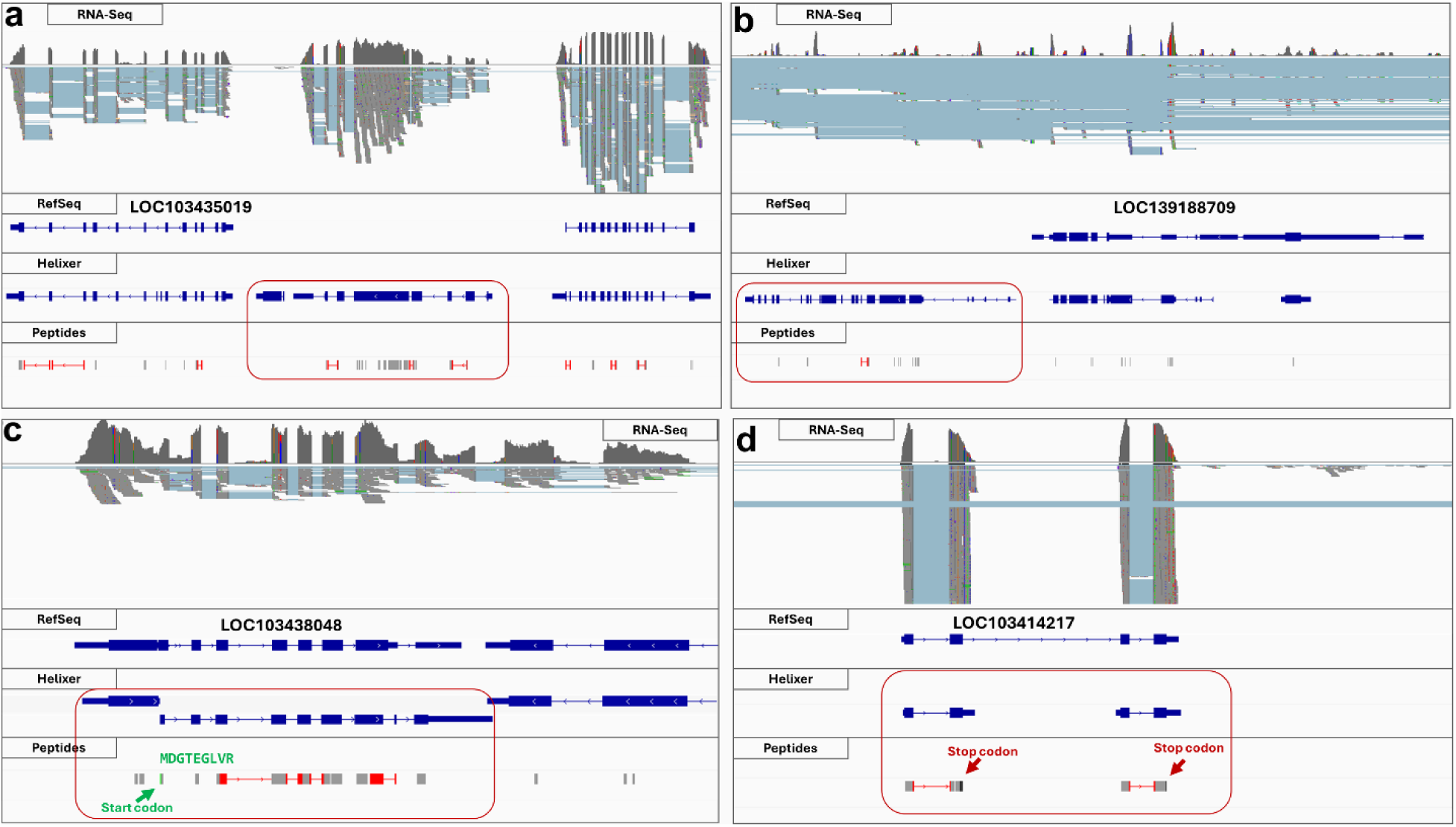
Proteogenomic validation and correction of RefSeq gene annotations in *Malus domestica*. Genome browser views showing loci where missing or mis-annotated RefSeq gene models are resolved using the GAP-MS proteogenomic pipeline. The top track displays RNA-seq read coverage. Gene models of RefSeq and Helixer annotations are shown in blue, mapped peptides in gray, and splice-junction peptides in red (bottom track). **(a)** GAP-MS recovers a functional gene absent from the reference RefSeq annotation, predicted by Helixer and supported by both peptide evidence and extensive RNA-seq coverage. **(b)** Identification of another gene missing from the NCBI RefSeq annotation: a TIR-NBS-LRR disease resistance protein containing an NB-ARC signaling domain required for pathogen recognition. **(c)** Correction of a falsely merged gene model. Helixer predicts two distinct adjacent gene models, whereas RefSeq incorrectly merges them into a single chimeric gene. GAP-MS identifies a peptide mapping precisely to the N-terminus of the downstream gene. The absence of an upstream tryptic cleavage site (R or K) confirms an independent translation start site. **(d)** Proteomic support for a gene split via C-terminal peptide mapping. An internal stop codon is confirmed by a C-terminal peptide lacking a subsequent tryptic cleavage site, corroborating the presence of two separate proteins rather than the merged RefSeq model.

A further example of a biological locus recovered by GAP-MS includes the TIR-NBS-LRR disease resistance protein (Fig. 6b). Accurate annotation of R-genes is often hindered by their characteristically low expression levels, which prevents RNA-seq datasets from providing sufficient evidence for automated gene prediction [53]. Furthermore, NB-LRR loci are frequently masked by annotation pipelines utilizing transposable element databases because these resistance genes are often misidentified as repetitive sequences [54]. Consequently, this specific gene is absent from the NCBI RefSeq annotation. GAP-MS identified proteomic support for the locus, including 13 internal peptides and one splice junction peptide. Domain analysis of the protein product confirmed a high prevalence of Leucine-Rich Repeats and NB-ARC signaling domains required for pathogen recognition. This discovery represents a broader trend across the dataset; systematic functional analysis of all novel gene models revealed a non-random distribution, significantly enriched for Gene Ontology (GO) terms associated with stress response and defense mechanisms (Supplementary Fig. 8).

Beyond discovering novel loci, GAP-MS demonstrates the capacity to correct structural errors in existing annotations, such as falsely merged genes. In one instance, the RefSeq annotation included a single continuous gene model "LOC103438048", whereas Helixer predicted two distinct, adjacent genes (Fig. 6c). GAP-MS provided definitive evidence supporting the split model by identifying a peptide mapping precisely to the N-terminus of the second predicted gene. The genomic sequence upstream of this peptide lacked the Arginine (R) or Lysine (K) residues required for tryptic cleavage, confirming the peptide represents a native biological N-terminus rather than a digestion artifact. This proves the existence of an independent translation initiation site in the middle of the RefSeq gene.

Structural refinement was also achieved for the gene model "LOC103414217", where C-terminal peptides supported a gene-split correctly predicted by Helixer but merged in RefSeq (Fig. 6d). While the stop codon at the end of the protein sequence was supported by a C-terminal peptide, another internal stop codon in the middle of the frame was confirmed by a second C-terminal peptide. Crucially, this internal peptide lacked a subsequent downstream tryptic cleavage site (not followed by R or K), providing physical evidence for an independent translation termination site. This confirms the Helixer prediction that the locus encodes two separate proteins rather than the single chimeric model presented in the standard reference.

## 5. Discussion

In this study, we introduce GAP-MS, a proteogenomic pipeline that integrates mass spectrometry evidence to evaluate and refine gene model predictions. By utilizing a high-quality subset from the forthcoming Crop Proteome Atlas, we applied this framework to 9 major crop species, demonstrating that integrating proteomic data allows us to confirm the translation of authentic coding sequences and point out possible prediction artifacts.

To evaluate the pipeline performance, we benchmarked GAP-MS against standard RefSeq annotations. While GAP-MS consistently improved precision across all tested tools, achieving substantial gains of up to 37%, it reduced recall when compared to reference annotations. However, standard reference annotations are not absolute ground truth and often contain errors such as pseudogenes, fragmented models, or transcriptionally silent regions. Thus, the observed drop in recall might reflect the correction of false positives in the reference rather than a loss of true genes.

The inherent limitations of mass spectrometry must also be considered when interpreting unsupported gene models. Proteomics is biased toward abundant proteins and is less sensitive to proteins with low expression or strong tissue- or condition-specificity. As a result, a genuine gene may be classified as unsupported simply because its protein product falls below the detection limit of the instrument or is not expressed in the sampled tissues. Increasing proteomic depth through strategies such as multi-protease digestion and extensive fractionation can partially address this limitation by improving sequence coverage and expanding the detectable proteome. Such workflows enable the detection of protein isoforms and sequence variants that are often missed by conventional shotgun approaches. Integrating these richer datasets into GAP-MS should therefore improve validation sensitivity and increase the number of novel gene models that can be supported [59].

Finally, GAP-MS was applied to identify novel coding regions, resulting in the recovery of functional genes missing from RefSeq (specifically those involved in stress response) showing its utility in improving genome annotation. Notably, across 9 crops more than 8,000 novel Helixer-predicted, peptide-supported gene models absent from current RefSeq annotations were identified. GAP-MS also demonstrated the capacity to correct structural errors in existing RefSeq reference annotations, such as resolving falsely merged chimeric models by identifying independent translation initiation and termination sites through specific N- and C-terminal peptides. To facilitate manual investigation, the pipeline also generates genomic coordinates of these mapped peptides in standard formats for seamless visualization in genome browsers.

Although not demonstrated in this work, GAP-MS could be extended with additional orthogonal evidence to improve sensitivity while preserving the distinction between inferred and directly observed proteins. Our ongoing work integrates transcriptomic data to provide evidence for transcript presence and to prioritize open reading frames that may be translationally repressed or fall below current proteomic detection limits. In parallel, advances in protein language models (PLMs) may help identify gene models with strong structural or evolutionary support despite lacking experimental protein evidence. Combining these orthogonal data types would create a more robust system that balances the strict specificity of proteomics with the broader sensitivity of transcriptomics and PLMs sequence analysis. Furthermore, de novo peptide sequencing offers a promising avenue to bypass the search space constraints of database-dependent proteogenomics. By deriving sequences directly from MS/MS spectra without a reference, this approach identifies translation products overlooked by initial gene prediction tools. Crucially, these de-novo-derived peptides can be seamlessly integrated into the GAP-MS input to facilitate the discovery of further novel genes.

Overall, GAP-MS represents a valuable advancement in the field of genome annotation, serving as an independent quality control filter. By validating gene models with physical evidence, the pipeline refines structural annotations and recovers biologically critical loci that are systematically missed by standard methods. The identification of overlooked genes, which are frequently fragmented or discarded due to their repetitive nature, highlights the immediate practical utility of this approach. In an era where accurate genomic resources are essential for modern breeding programs, the systemic loss of such agronomic traits in reference annotations compromises crop improvement strategies. Therefore, as the demand for high-quality functional genomics grows, we are convinced that integrating direct proteomic evidence through pipelines like GAP-MS will be essential for defining high-confidence reference proteomes and unlocking the full functional potential of crop genomes.

## 6. Data Availability

All computational scripts and datasets generated in this study are available on GitHub: https://github.com/qussai96/GAP-MS.

Proteomics data are available via ProteomeXchange with identifier **PXD076526**

Identified peptides from mass spectrometry data of 9 crops together with gene models predicted with Braker2, Galba, Helixer and Annevo are available on Zenodo: **10.5281/zenodo.18445783**.

## 7. Supplementary Data statement

Supplementary Data are available at *NAR* Online.

## 8. Authors’ contributions

Q.A., B.K. and D.F. jointly conceived the study. Q.A. wrote the main manuscript text, developed the pipeline, conducted the benchmarking of the tools, and created all plots and figures. D.F., M.W. and B.K. acquired the financial support for this project, conceptualized and supervised the work, in addition to reviewing and refining the manuscript. M.W. supervised the proteomics analyses, reviewed and edited the manuscript. B.K. supported the project with his expertise in proteomics and contributed to the revision process. All authors read and approved the final manuscript.

## 10. Acknowledgements

The authors are grateful to all members of the ‘Proteomes that Feed the World’ international program for their essential contributions to the cultivation, sample preparation, and mass spectrometry data acquisition for the 9 crop species utilized in this study. Their foundational work is part of a coordinated effort to establish the Crop Proteome Atlas.

## 13. Supplementary figures

**Supplementary Fig 1.**
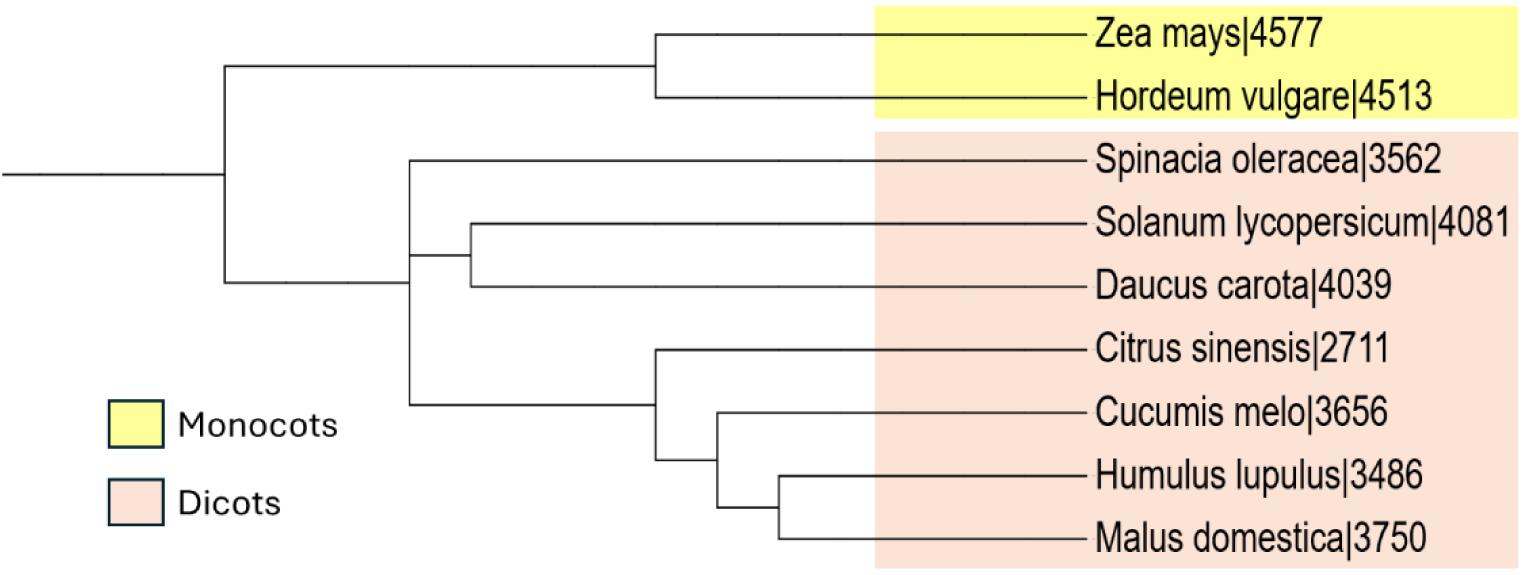
Phylogenetic tree of the selected 9 crop species (Species | TaxID). Selected species span diverse taxonomic groups and plant classes (monocots and dicots). The tree was visualized and annotated using the interactive Tree of Life (iTOL) tool using NCBI taxonomy IDs.

**Supplementary Fig 2.**
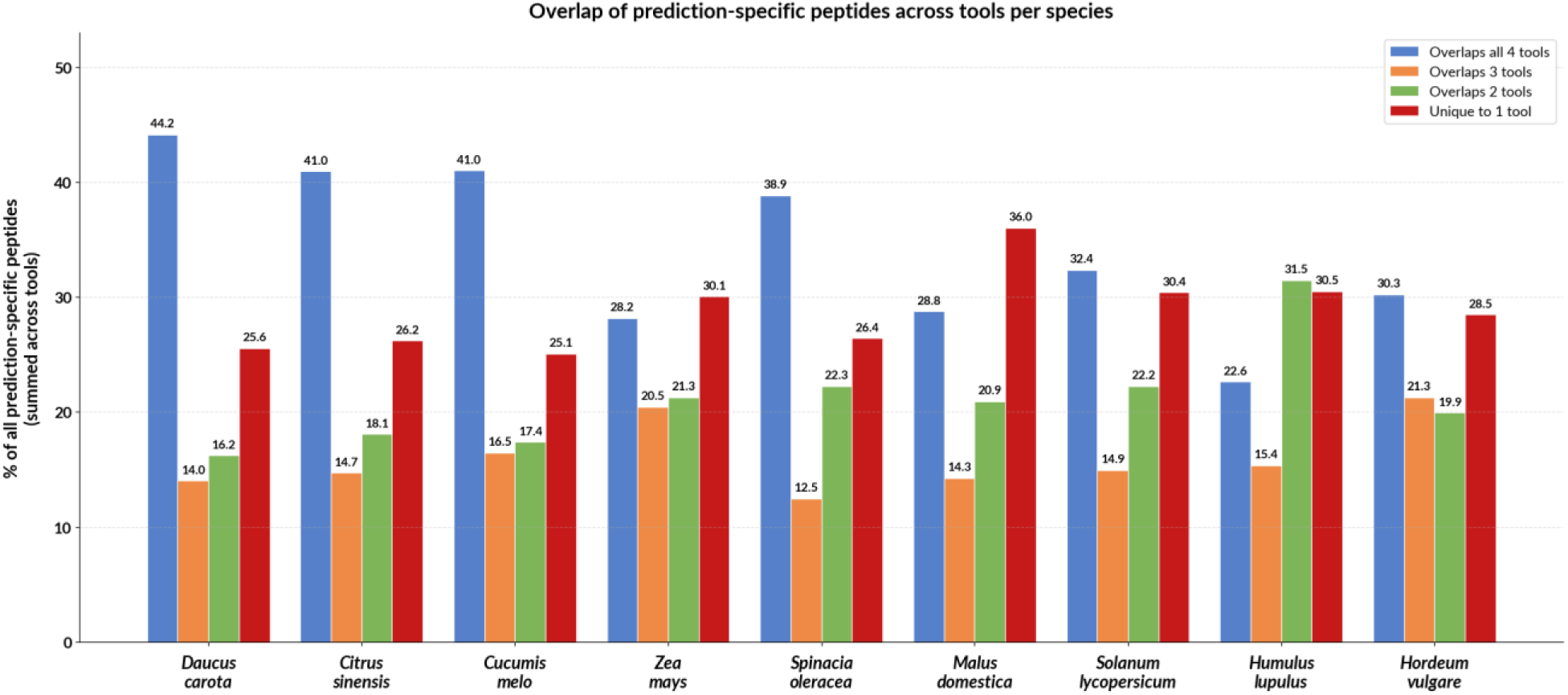
Overlap of prediction-specific peptides between the four gene prediction tools. Grouped bar plot showing the percentage of prediction-specific peptides (potential novel coding sequences absent from the RefSeq annotation) per species according to their overlap across the four gene prediction tools. Bars indicate peptides detected by all four tools, by three tools, by two tools, or uniquely by one tool.

**Supplementary Fig 3.**
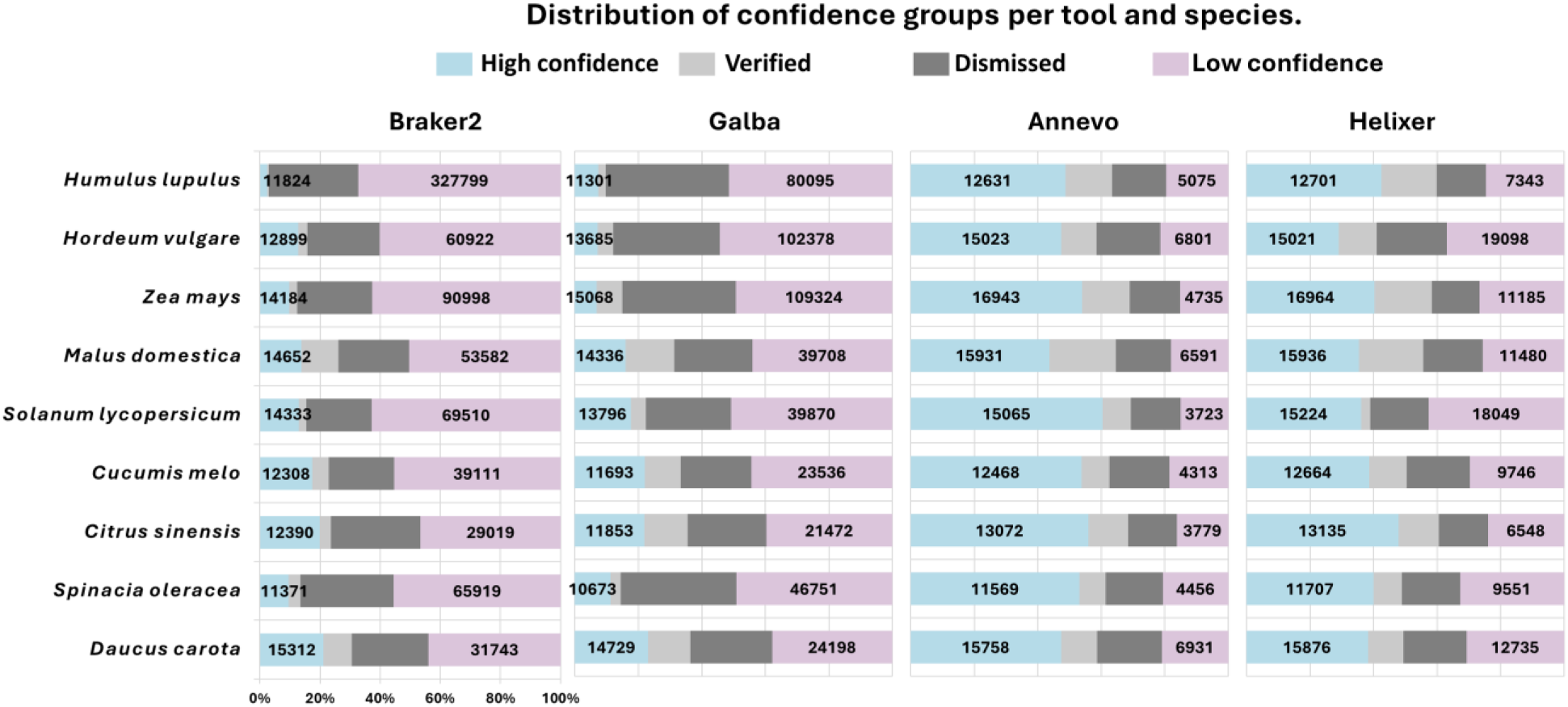
Distribution of confidence groups per species. Stacked bars show the proportion of proteins assigned to high-confidence (blue) and low-confidence (pink) training sets, as well as the unlabeled proteins subsequently classified as supported (light gray) or unsupported (dark gray) across 9 crops and 4 prediction tools.

**Supplementary Fig 4.**
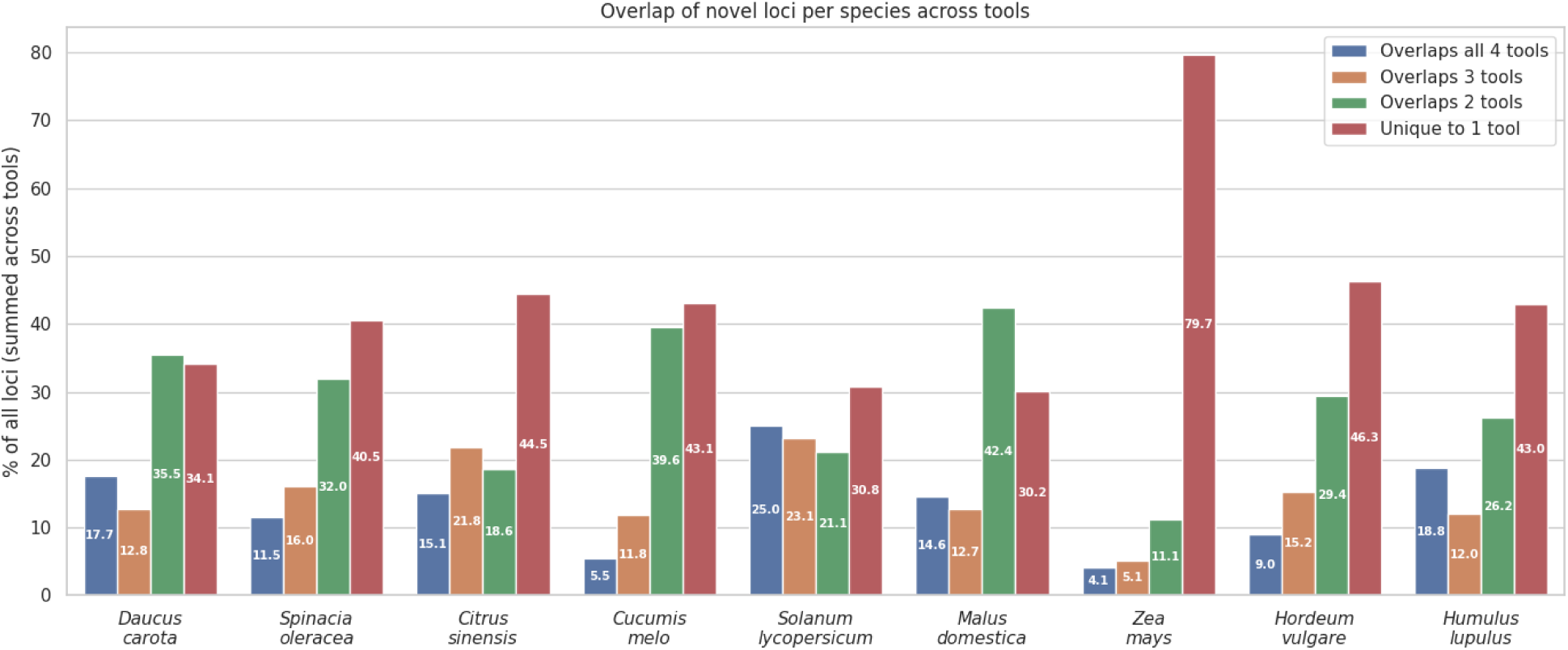
Overlap of novel peptide-supported loci across prediction tools. Grouped bar plot showing the percentage of novel peptide-supported loci per species according to their overlap across the four gene prediction tools. Bars indicate loci detected by all four tools, by three tools, by two tools, or uniquely by one tool.

**Supplementary Fig 5.**
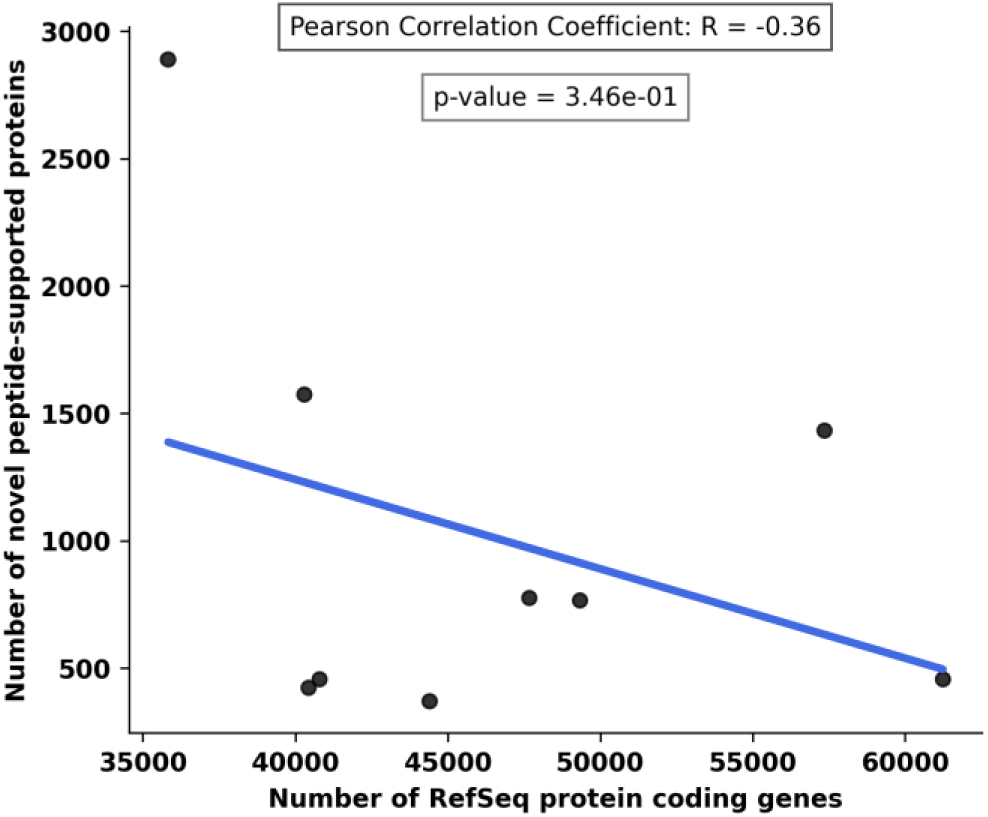
Correlation between the number of RefSeq protein coding genes and the number of novel peptide-supported Helixer genes. A p-value of 0.34 indicated no significant correlation.

**Supplementary Fig 6.**
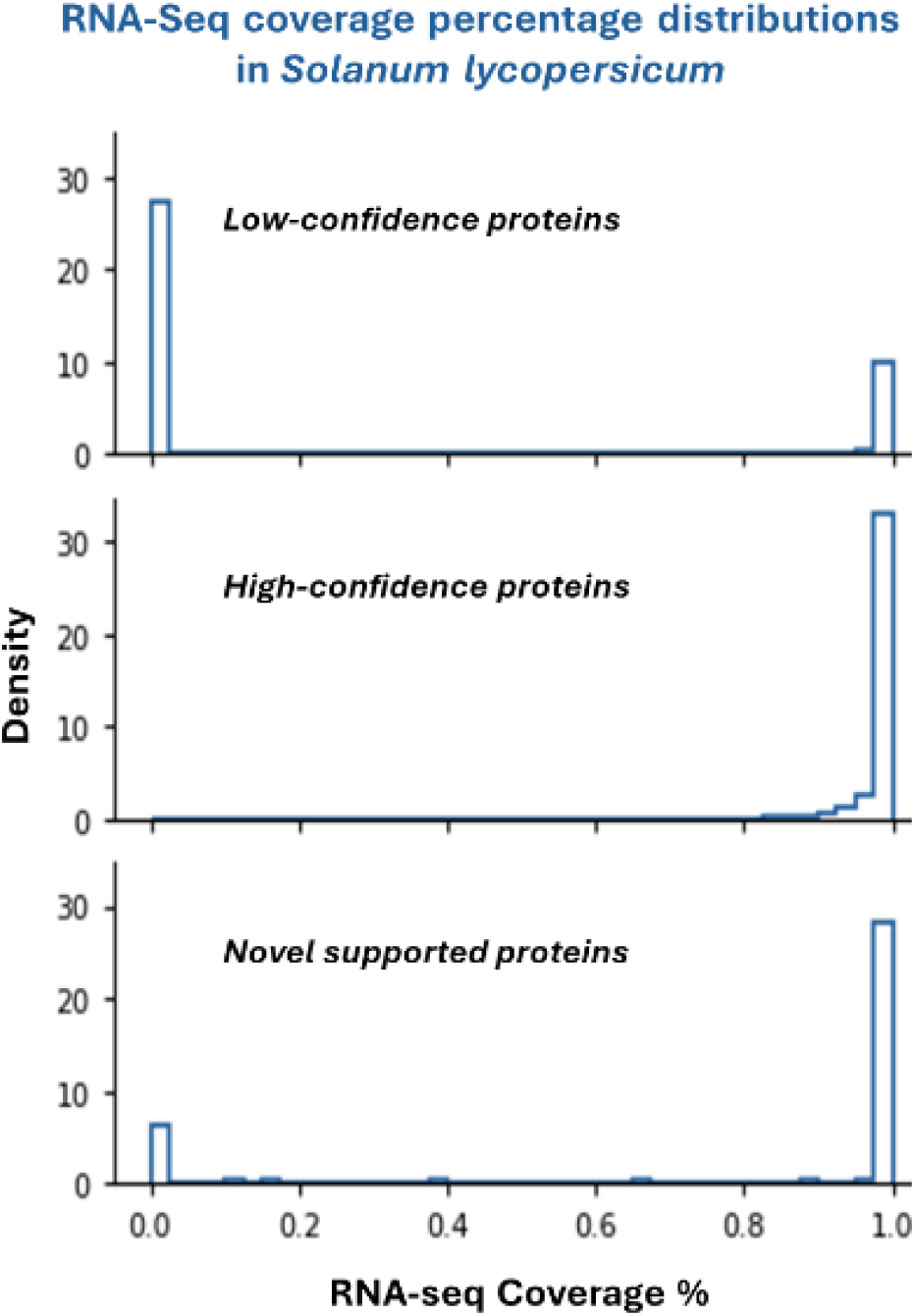
Density distributions of RNA-seq coverage percentages in *Solanum lycopersicum* (tomato). The panels compare the coverage profiles of high-confidence reference proteins (top), low-confidence reference proteins (middle), and the novel peptide-supported proteins (bottom), highlighting the similarity in distribution between the novel and high-confidence sets.

**Supplementary Fig 7.**
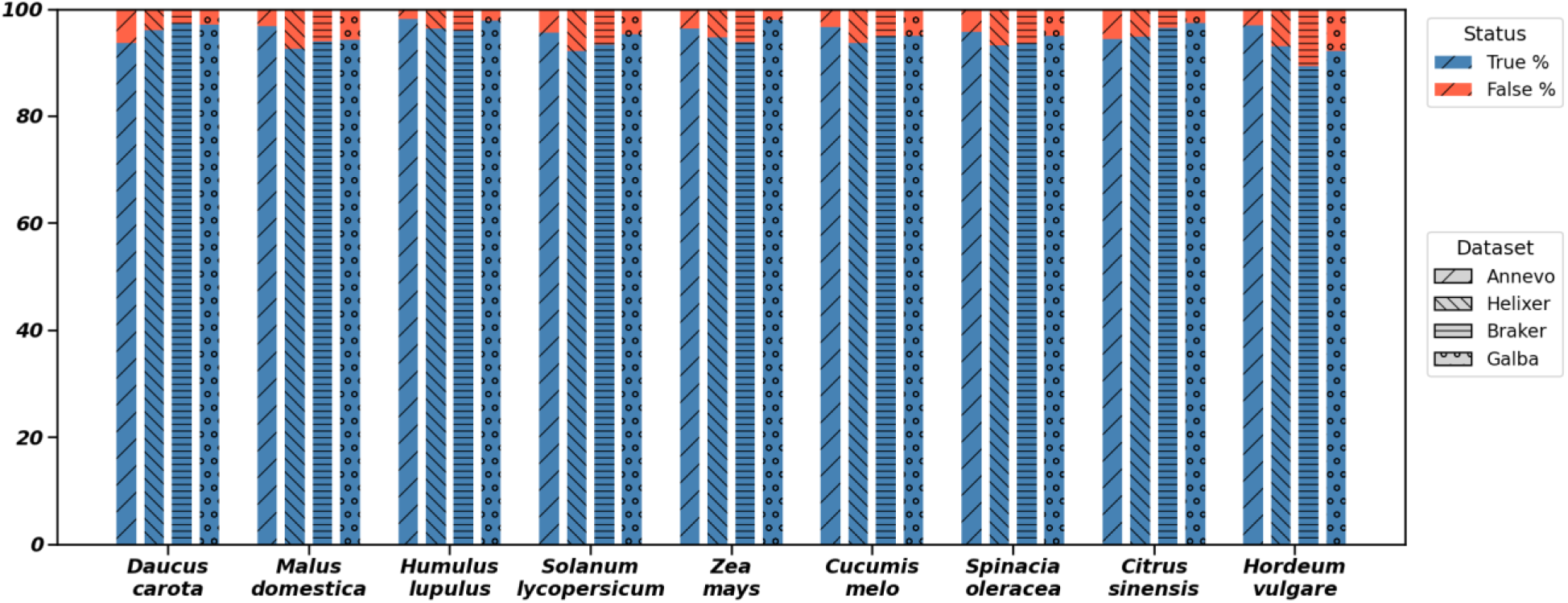
Quality assessment of novel peptide-supported proteins. Proportion of novel Helixer-predicted proteins (supported by GAP-MS but absent from RefSeq) that were classified as valid coding sequences (True) versus invalid (False) by Psauron.

**Supplementary Fig 8.**
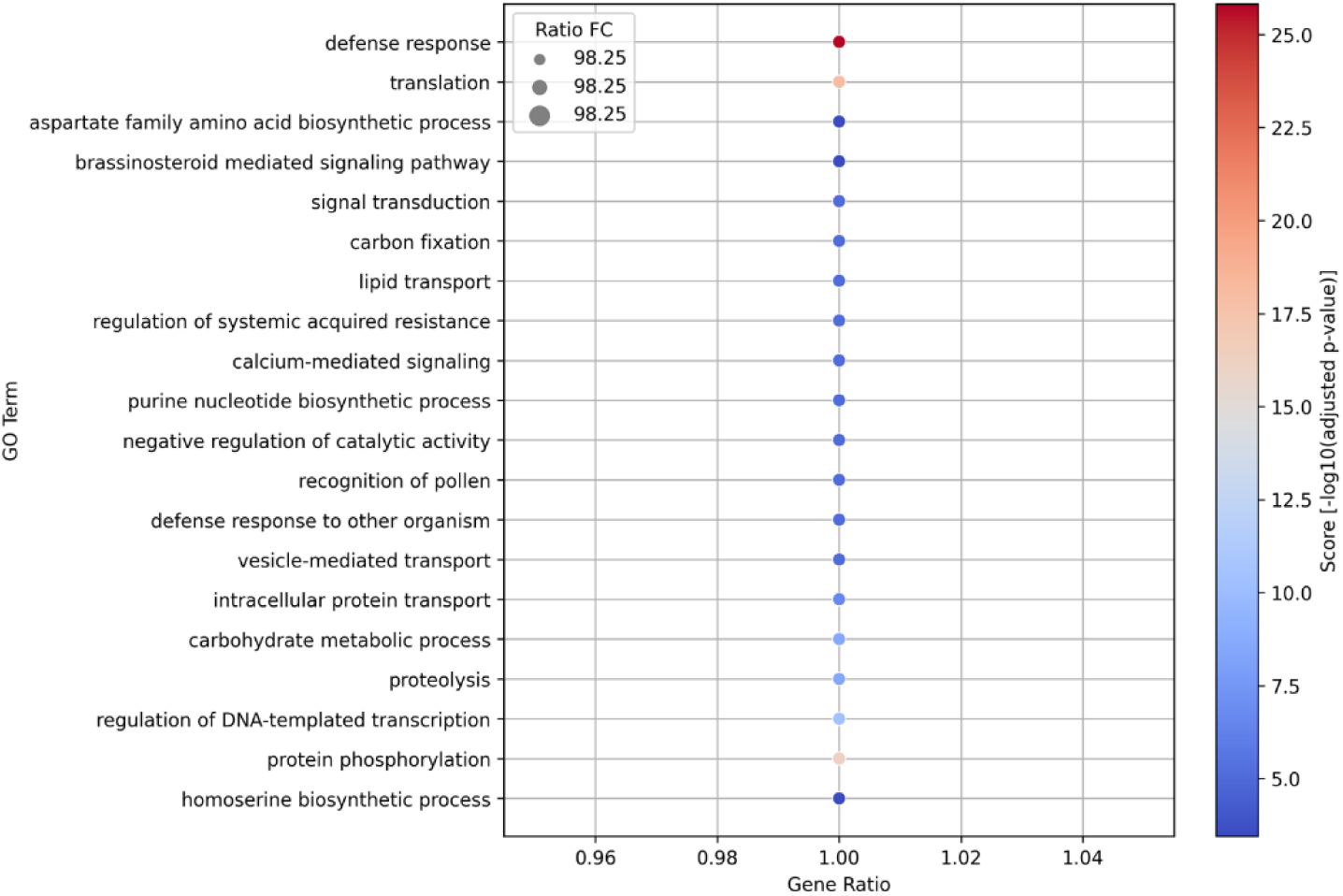
GO enrichment analysis of novel helixer peptide supported genes in *Malus domestica*. The y-axis lists enriched GO biological processes, showing that defense response emerging as the top enriched process, which indicated that the gene set is dominated by stress- or immunity-related functions

