## Supplementary Table 1 for "GAP-MS: Automated validation of gene predictions using integrated mass spectrometry evidence"

| Accession | Potanical_Name | Common_Name | Assembly_Level |
|---|---|---|---|
| GCF_000003195.3 | Sorghum bicolor | sorghum | Chromosome |
| GCF_000004515.6 | Glycine max | soybean | Chromosome |
| GCF_000346465.2 | Prunus persica | peach | Chromosome |
| GCA_963454935.2 | Ilex paraguariensis | Yerba Mate | Scaffold |
| GCF_904849725.1 | Hordeum vulgare | barley | Chromosome |
| GCF_963169125.1 | Humulus lupulus | European hop | Chromosome |
| GCF_009730915.1 | Dioscorea cayenensis | Guinea yam | Chromosome |
| GCF_002870075.4 | Lactuca sativa | garden lettuce | Chromosome |
| GCA_048183575.1 | Vicia faba | fava bean | Chromosome |
| GCA_903112645.1 | Prunus armeniaca | apricot | Contig |
| GCF_030704535.1 | Vitis vinifera | wine grape | Complete Genome |
| GCF_036785885.1 | Coffea arabica | coffee | Chromosome |
| GCF_042453785.1 | Malus domestica | apple | Complete Genome |
| GCF_025177605.1 | Cucumis melo | muskmelon | Chromosome |
| GCF_003086295.3 | Arachis hypogaea | peanut | Chromosome |
| GCF_000004075.3 | Cucumis sativus | cucumber | Chromosome |
| GCF_026745355.1 | Beta vulgaris | Sugar beet | Chromosome |
| GCF_963583255.1 | Pyrus communis | pear | Chromosome |
| GCF_002878395.1 | Capsicum annuum | Green pepper | Chromosome |
| GCF_000442705.2 | Elaeis guineensis | African oil palm | Chromosome |
| GCF_001683475.1 | Chenopodium quinoa | quinoa | Scaffold |
| GCA_015342445.1 | Digitaria exilis | white fonio | Contig |
| GCF_001531365.2 | Cynara cardunculus | artichoke | Chromosome |
| GCA_051132755.1 | Persea americana | avocado | Complete Genome |
| GCF_902167145.1 | Zea mays | Maize | Chromosome |
| GCF_000499845.2 | Phaseolus vulgaris | common bean | Chromosome |
| GCF_024323335.1 | Pisum sativum | garden pea | Chromosome |
| GCF_036512215.1 | Solanum lycopersicum | tomato | Chromosome |
| GCF_004118075.2 | Vigna unguiculata | cowpea | Chromosome |
| GCF_000695525.1 | Brassica oleracea | cabbage | Chromosome |
| GCF_001659605.2 | Manihot esculenta | cassava | Chromosome |
| GCF_000331145.2 | Cicer arietinum | chickpea | Chromosome |
| GCF_002127325.2 | Helianthus annuus | common sunflower | Chromosome |
| GCF_009389715.1 | Phoenix dactylifera | date palm | Chromosome |
| GCF_018446385.1 | Zingiber officinale | Ginger | Chromosome |
| GCF_020379485.1 | Brassica napus | rape | Chromosome |
| GCF_020520425.1 | Spinacia oleracea | spinach | Chromosome |
| GCF_000226075.1 | Solanum tuberosum | potato | Scaffold |
| GCF_022201045.2 | Citrus sinensis | sweet orange | Complete Genome |
| GCF_901000735.1 | Corylus avellana | Hazelnut | Chromosome |
| GCF_001876935.1 | Asparagus officinalis | garden asparagus | Chromosome |
| GCF_000512975.1 | Sesamum indicum | sesame | Chromosome |
| GCF_001411555.2 | Juglans regia | Persian walnut | Chromosome |
| GCF_000340665.2 | Cajanus cajan | pigeon pea | Chromosome |
| GCA_039639745.1 | Bertholletia excelsa | Brazil nut | Chromosome |
| GCF_002806865.2 | Cucurbita pepo | vegetable marrow | Chromosome |
| GCF_002742605.1 | Olea europaea | Olive | Chromosome |

| | | | |
|---|---|---|---|
| GCF_034140825.1 | Oryza sativa | Japanese rice | Complete Genome |
| GCF_018294505.1 | Triticum aestivum | bread wheat | Chromosome |
| GCF_003573695.1 | Papaver somniferum | opium poppy | Chromosome |
| GCF_011075055.1 | Mangifera indica | mango | Chromosome |
| GCF_029168945.1 | Cannabis sativa | hemp | Chromosome |
| GCA_004153795.1 | Camellia sinensis | Tea | Scaffold |
| GCF_002207925.1 | Prunus avium | sweet cherry | Scaffold |
| GCF_008641045.1 | Pistacia vera | Pistachio | Scaffold |
| GCF_001540865.1 | Ananas comosus | pineapple | Chromosome |
| GCF_001625215.2 | Daucus carota | carrot | Chromosome |
| GCF_902201215.1 | Prunus dulcis | almond | Chromosome |
| GCA_021397845.1 | Areca catechu | Areca catechu (betel pal | Chromosome |
| GCA_022606695.1 | Vaccinium macrocarpon | Vaccinium macrocarpor | Chromosome |
| GCA_015708375.1 | Cydonia oblonga | Cydonia oblonga (quince | Contig |
| GCA_014218235.1 | Colocasia esculenta | Colocasia esculenta (tar | Chromosome |
| GCA_008629595.1 | Brassica rapa | Brassica rapa subsp. pel | Chromosome |
| GCA_042847195.1 | Ficus carica | Ficus carica (common fi | Chromosome |
| GCA_034638355.1 | Brassica oleracea | Brassica oleracea var. ca | Chromosome |
| GCA_041146485.1 | Mentha x piperita | Mentha x piperita (peppe | Chromosome |
| GCA_012066045.3 | Morus alba | Morus alba (white mulbe | Chromosome |
| GCA_051167515.1 | Linum usitatissimum | Linum usitatissimum (fla | Chromosome |
| GCA_034640245.1 | Brassica oleracea | Brassica oleracea var. b | Chromosome |
| GCA_026122585.1 | Anacardium occidentale | Anacardium occidentale | Scaffold |
| GCA_036245055.1 | Diospyros kaki | Diospyros kaki (kaki pers | Contig |
| GCF_040712315.1 | Castanea sativa | Castanea sativa (Europe | Chromosome |
| GCA_001633085.1 | Carthamus tinctorius | Carthamus tinctorius (sa | Scaffold |
| GCA_019916065.1 | Vitellaria paradoxa | Vitellaria paradoxa (shea | Chromosome |
| GCA_051171125.1 | Avena sativa | Avena sativa (oats) | Chromosome |
| GCA_047496405.1 | Rubus idaeus | Rubus idaeus (red raspb | Scaffold |
| GCA_023846275.1 | Vanilla planifolia | Vanilla planifolia (cultiva | Chromosome |
| GCA_033239045.1 | Fagopyrum esculentum | Fagopyrum esculentum | Scaffold |
| GCA_014504835.1 | Vaccinium corymbosum | Vaccinium corymbosum | Scaffold |
| GCA_030737875.1 | Allium sativum | Allium sativum (garlic) | Chromosome |
| GCA_030765085.1 | Allium cepa | Allium cepa (onion) | Chromosome |
| GCA_965641915.1 | Secale cereale | Secale cereale (rye) | Chromosome |
| GCA_034509205.1 | Ceratonia siliqua | Ceratonia siliqua (carob | Scaffold |
| GCA_018398585.1 | Simmondsia chinensis | Simmondsia chinensis (j | Scaffold |
| GCA_965211865.1 | Ribes uva-crispa | Ribes uva-crispa (Englisl | Chromosome |
| GCA_963924085.1 | Cenchrus americanus | Cenchrus americanus (p | Chromosome |
| GCA_020631735.1 | Saccharum officinarum | Saccharum officinarum | Chromosome |
| GCA_002525835.2 | Ipomoea batatas | Ipomoea batatas (sweet | Chromosome |
| GCA_034663555.1 | Citrus x limon | Citrus x limon (lemon) | Chromosome |
| GCA_034640255.1 | Brassica oleracea | Brassica oleracea var. ita | Chromosome |
| GCA_034370585.1 | Fragaria x ananassa | Fragaria x ananassa (stra | Complete Genome |
| GCA_036169575.1 | Citrus reticulata | Citrus reticulata (manda | Chromosome |
| GCA_024500025.1 | Syzygium aromaticum | Syzygium aromaticum (c | Chromosome |
| GCA_965178245.1 | Ribes rubrum | Ribes rubrum (red curra | Chromosome |
| GCA_003604295.2 | Cocos nucifera | Cocos nucifera (coconut | Chromosome |

| | | | |
|---|---|---|---|
| GCA_039639715.1 | Citrullus lanatus | Citrullus lanatus (waterr | Complete Genome |
| GCA_051167385.1 | Actinidia chinensis | Actinidia chinensis var. ( | Chromosome |
| GCA_018132145.1 | Boehmeria nivea | Boehmeria nivea var. ter | Chromosome |
| GCA_000340665.3 | Cajanus cajan | Cajanus cajan (pigeon p | Chromosome |
| GCA_035048815.1 | Abelmoschus esculentus | Abelmoschus esculentu | Scaffold |
| GCA_018703725.1 | Brassica juncea | Brassica juncea (brown ı | Chromosome |
| GCA_003724115.2 | Anethum foeniculum | Anethum foeniculum (fe | Scaffold |
| GCF_000150535.2 | Carica papaya | papaya | Scaffold |
| GCF_000208745.1 | Theobroma cacao | cacao | Chromosome |

| Assembly_Sequencing_Tech | Assembly_Assembly_Me | mbly_Release_Date | Total_Sequence_Leng |
|---|---|---|---|
| Sanger; Illumina | ARACHNE_modified v. 20( | 07-04-17 | 708735318 |
| ABI 3739 | ARACHNE v. 200721016_i | 10-03-21 | 978386919 |
| ABI 3739 | ARACHNE v. 20071016_m | 02-02-17 | 227411381 |
| Illumina, PacBio | Quickmerg | 04-03-24 | 1064802823 |
| PacBio RSII | Hi-Canu (commit r9818) | 01-04-21 | 4225577519 |
| PacBio,Arima2 | various | 19-08-23 | 2488098718 |
| Nanopore PromethION | flye v. 2.4.2; pilon v. 1.23; | 25-11-19 | 584153202 |
| PacBio Sequel II | Hifiasm v. Dec-2011; BioN | 23-11-22 | 2589853228 |
| Pacbio Sequel II | hifiasm v. 0.16.1 | 28-02-25 | 11807183301 |
| PacBio Sequel | flye(v2.6) | 08-07-20 | 215952222 |
| PacBio Sequel | hifiasm v. v.15 | 09-08-23 | 494873210 |
| Illumina NextSeq; PacBio Sequel | HiFiasm v. 2023-01-20 | 23-02-24 | 1198133039 |
| PacBio RSII | HiFiasm v. v0.19.5-r587 | 27-09-24 | 650775246 |
| PacBio HiFi Sequel II | HifiASM v. 0.5 | 14-09-22 | 438252206 |
| PacBio; HiC | MECAT v. 2017-07; Arrow | 16-10-23 | 2557413415 |
| PacBio RSII; PacBio Sequel; 10X Ge | Celera Assembler v. 1.8; F | 15-11-19 | 224801081 |
| PacBio RSII | FALCON v. 0.2.2 | 09-12-22 | 568751015 |
| PacBio,Arima2 | various | 05-11-23 | 487337232 |
| Illumina HiSeq | Supernova v. 1.1-Chili-Pe | 30-01-18 | 3211819707 |
| PacBio RSII | FALCON via SMRT-Analysi | 09-08-24 | 1841922575 |
| PacBio | smrtmake v. 1.0; IrysView | 19-05-17 | 1333398936 |
| PacBio | Canu v. 1.8 | 10-11-20 | 760650579 |
| Illumina HiSeq | AllPaths v. 41684 | 17-04-18 | 724962400 |
| PacBio RSII; Oxford Nanopore Prom | hifiasm v. 0.19.5-r587 | 28-06-25 | 841551734 |
| PacBio Sequel | Canu 1.8 | 21-03-20 | 2182075994 |
| PacBio | FALCON v. unknown | 23-04-24 | 537154354 |
| PacBio Sequel | Canu v. 1.8; Pilon v. 1.23; | 19-07-22 | 3796066963 |
| PacBio Sequel II | Hifiasm v. 0.16.1 | 03-09-24 | 832773323 |
| PacBio; Bionano | CANU-Abruijn-FALCON v. | 30-01-19 | 518595462 |
| Illumina GAII; Illumina HiSeq; 454 | SOAPdenovo v. 1.05 | 27-05-14 | 488593889 |
| PacBio RSII; Sequel | Canu v. 1.6 | 13-08-21 | 639586700 |
| Illumina HiSeq | 3D-DNA v. 180922 | 16-12-24 | 530271971 |
| PacBio RSII | CANU v. 1.3; CANU v. 1.4; | 15-07-20 | 3009595538 |
| PacBio | Falcon v. 2017.06.28-18.0 | 04-11-19 | 772315838 |
| Oxford Nanopore PromethION | SMARTDENOVO v. 1 | 21-05-21 | 3090266393 |
| PacBio Sequel; Illumina HiSeq | Canu v. 1.6; Pilon v. 1.22; | 08-10-21 | 1001499700 |
| PacBio Sequel | Canu v. 1.7.1 | 20-10-21 | 894256318 |
| Illumina GA2 | SOAPdenovo v. 1014 | 19-09-11 | 705779115 |
| PacBio Sequel II | FALCON-Unzip v. 2.0; CAN | 11-02-22 | 298977511 |
| Illumina,Oxford Nanopore | MaSuRCA-3.2.6,HiRise | 25-11-20 | 369779143 |
| Illumina HiSeq | SOAPdenovo2 v. r240; PBj | 06-02-17 | 1187538024 |
| Illumina Hiseq 2000 | SOAPdenovo v. 1.06 | 06-01-14 | 274906174 |
| Illumina HiSeq; Oxford Nanopore M | MaSuRCA v. 3.2.1; HiRise | 08-07-20 | 572786133 |
| Illumina Hiseq2000 | SOAPdenovo v. 1.05 | 29-09-16 | 590369945 |
| PacBio Sequel | Canu v. 1.8; HiRise v. MAY | 14-05-24 | 599383329 |
| Illumina HiSeq | SOAPdenovo v. r240; Brea | 07-12-17 | 259381069 |
| Illumina HiSeq | SOAPdenovo v. 2 | 03-11-17 | 1140989389 |

| | | | |
|---|---|---|---|
| PacBio Sequel II; Nanopore GridION | Hifiasm v. 0.19.3-r572 | 06-12-23 | 385710679 |
| PacBio; Illumina HiSeq | FALCON v. 0.2.2; DenovoM | 06-05-21 | 14566502436 |
| Illumina HiSeq; PacBio; 10X Genom | DeNovoMAGIC v. 3.0; Falc | 18-09-18 | 2715377404 |
| Pacbio Sequel II | Canu v. 1.8 | 10-03-20 | 391961648 |
| Oxford Nanopore GridION; PacBio F | NextDenovo v. 2.3.1; SMA | 14-03-23 | 770269637 |
| Illumina Hiseq 2500; PacBio RSII | Platanus v. 1.24; SOAPder | 11-02-19 | 3105212962 |
| Illumina HiSeq 2000 | SOAPdenovo v. 2-rev240; | 12-06-17 | 272361615 |
| PacBio Sequel | PBJelly v. 15.8.24 | 23-09-19 | 671119336 |
| Illumina HiSeq; PacBio; 454 | AllPaths v. OCT-2012 | 14-03-16 | 381896302 |
| PacBio Sequel; Oxford Nanopore; Il | FALCON v. 0.3.0; CANU v. | 20-10-23 | 440713325 |
| Illumina, ONT | MASURCA,Redundans,SS | 10-10-19 | 227599157 |
| NA | NA | 11-01-22 | 2823079964 |
| NA | NA | 18-03-22 | 484919498 |
| NA | NA | 01-12-20 | 488422409 |
| NA | NA | 17-08-20 | 2405846852 |
| NA | NA | 20-09-19 | 234688483 |
| NA | NA | 09-10-24 | 318315163 |
| NA | NA | 28-12-23 | 570744922 |
| NA | NA | 09-08-24 | 2040613434 |
| NA | NA | 07-01-21 | 336455645 |
| NA | NA | 01-07-25 | 494853525 |
| NA | NA | 28-12-23 | 534419652 |
| NA | NA | 10-11-22 | 356592713 |
| NA | NA | 29-01-24 | 708766281 |
| NA | NA | 15-07-24 | 716029944 |
| NA | NA | 27-04-16 | 661937708 |
| NA | NA | 09-09-21 | 667213145 |
| NA | NA | 02-07-25 | 10935490949 |
| NA | NA | 06-02-25 | 276910634 |
| NA | NA | 01-08-22 | 1416355720 |
| NA | NA | 02-11-23 | 1266838060 |
| NA | NA | 09-09-20 | 393150603 |
| NA | NA | 16-08-23 | 16462794857 |
| NA | NA | 17-08-23 | 15932385976 |
| NA | NA | 19-07-25 | 6702647067 |
| NA | NA | 21-12-23 | 477340296 |
| NA | NA | 19-05-21 | 831537837 |
| NA | NA | 17-03-25 | 813097327 |
| NA | NA | 09-12-24 | 1914840079 |
| NA | NA | 27-10-21 | 6804885551 |
| NA | NA | 16-10-17 | 837013208 |
| NA | NA | 28-12-23 | 682876182 |
| NA | NA | 28-12-23 | 557942728 |
| NA | NA | 14-12-23 | 793463571 |
| NA | NA | 26-01-24 | 303165851 |
| NA | NA | 03-08-22 | 370257752 |
| NA | NA | 11-03-25 | 834940214 |
| NA | NA | 28-08-24 | 2687377809 |

| | | | |
|---|---|---|---|
| NA | NA | 14-05-24 | 373729396 |
| NA | NA | 01-07-25 | 607462286 |
| NA | NA | 26-04-21 | 270213153 |
| NA | NA | 20-12-24 | 594702767 |
| NA | NA | 03-01-24 | 1194595883 |
| NA | NA | 04-06-21 | 933495403 |
| NA | NA | 24-09-21 | 1011385054 |
| | | 06-05-08 | 369781828 |
| Roche/454 GSFLX, Illumina HiSeq, | Newbler, SSPACE, ScaffR[illegible] | 09-07-16 | 324719311 |

| Total_Ungapped_Leng | Total_gaps_percen | Number_of_Conti | Number_of_Scaffolds | Number_o | Number_o |
|---|---|---|---|---|---|
| 675363888 | 4.71 | 4773 | 867 | 10 | 2 |
| 952478023 | 2.65 | 9197 | 345 | 20 | 2 |
| 224638928 | 1.22 | 2530 | 191 | 8 | 1 |
| 1043925183 | 1.96 | 24189 | 10611 | | |
| 4224251725 | 0.03 | 452 | 290 | 7 | 1 |
| 2488073467 | 0.00 | 448 | 319 | 10 | |
| 579406249 | 0.81 | 7012 | 2253 | 28 | 1 |
| 2588515351 | 0.05 | 475 | 91 | 9 | 2 |
| 11806989281 | 0.00 | 3099 | 1158 | 7 | |
| 215952222 | 0.00 | 93 | 93 | | |
| 494873210 | 0.00 | 19 | 19 | 19 | 2 |
| 1198080103 | 0.00 | 239 | 133 | 22 | 1 |
| 650775246 | 0.00 | 17 | 17 | 17 | 2 |
| 438228306 | 0.01 | 1548 | 1309 | 12 | 1 |
| 2553632056 | 0.15 | 4139 | 442 | 20 | 1 |
| 224764911 | 0.02 | 173 | 84 | 7 | 4 |
| 534762357 | 5.98 | 3098 | 18 | 9 | 2 |
| 487308232 | 0.01 | 176 | 31 | 17 | |
| 3124034419 | 2.73 | 133777 | 81200 | 12 | 2 |
| 1714158544 | 6.94 | 37553 | 29 | 16 | 1 |
| 1322160028 | 0.84 | 4211 | 3486 | | 1 |
| 760650579 | 0.00 | 3329 | 3329 | | |
| 654470093 | 9.72 | 73303 | 13479 | 17 | |
| 841551734 | 0.00 | 12 | 12 | 12 | |
| 2178268108 | 0.17 | 1393 | 685 | 10 | 2 |
| 531494354 | 1.05 | 1032 | 1032 | 11 | 2 |
| 3785824465 | 0.27 | 2392 | 1566 | 7 | |
| 832715554 | 0.01 | 178 | 12 | 12 | 2 |
| 515874664 | 0.52 | 743 | 675 | 11 | 1 |
| 445594160 | 8.80 | 83955 | 32885 | 9 | 1 |
| 637114355 | 0.39 | 756 | 102 | 18 | 2 |
| 482665111 | 8.98 | 39952 | 6445 | 8 | 1 |
| 2947692501 | 2.06 | 2712 | 332 | 17 | 2 |
| 772284138 | 0.00 | 2706 | 2389 | 18 | 2 |
| 3090199650 | 0.00 | 1108 | 210 | 22 | 1 |
| 1001413903 | 0.01 | 4004 | 3164 | 19 | 3 |
| 894250018 | 0.00 | 307 | 244 | 6 | 2 |
| 663134508 | 6.04 | 60068 | 14853 | | 1 |
| 298977511 | 0.00 | 9 | 9 | 9 | 2 |
| 368016482 | 0.48 | 14543 | 11 | 11 | 1 |
| 1152016560 | 2.99 | 56182 | 11792 | 10 | |
| 270333346 | 1.66 | 26175 | 16368 | 16 | 1 |
| 572182264 | 0.11 | 25807 | 24756 | 16 | 1 |
| 555945622 | 5.83 | 67833 | 31510 | 11 | 1 |
| 599348329 | 0.01 | 367 | 17 | 17 | |
| 244760851 | 5.64 | 30570 | 24193 | 20 | 1 |
| 1031505450 | 9.60 | 83943 | 41225 | 23 | 1 |

| | | | | | |
|---|---|---|---|---|---|
| 385710679 | 0.00 | 12 | 12 | 12 | 3 |
| 14314044360 | 1.73 | 306348 | 91588 | 21 | 1 |
| 2709971134 | 0.20 | 65343 | 34380 | 11 | 1 |
| 391878804 | 0.02 | 419 | 250 | 20 | |
| 770264337 | 0.00 | 70 | 17 | 10 | 1 |
| 2860303599 | 7.89 | 93350 | 14027 | | 1 |
| 246800503 | 9.38 | 32310 | 10148 | | |
| 670060100 | 0.16 | 2326 | 1864 | | 1 |
| 375114483 | 1.78 | 9390 | 3128 | 25 | 1 |
| 440657925 | 0.01 | 563 | 9 | 9 | 2 |
| 223688024 | 1.72 | 4395 | 691 | 8 | 1 |
| 2778792366 | 1.568768812 | 7480 | 57 | 16 | 0 |
| 484908398 | 0.00228904 | 124 | 1 | 12 | 0 |
| 488422409 | 0 | 303932 | 303932 | 0 | 0 |
| 2405007904 | 0.034871214 | 16127 | 7721 | 14 | 0 |
| 232151983 | 1.080794408 | 28786 | 3411 | 10 | 0 |
| 316264485 | 0.644228814 | 264 | 0 | 13 | 0 |
| 570679922 | 0.011388625 | 159 | 20 | 9 | 0 |
| 2040596634 | 0.000823282 | 762 | 522 | 72 | 0 |
| 336321135 | 0.039978524 | 519 | 234 | 14 | 1 |
| 494848525 | 0.0010104 | 20 | 0 | 15 | 0 |
| 534419652 | 0 | 72 | 63 | 9 | 0 |
| 354997678 | 0.447298821 | 5216 | 3268 | 0 | 0 |
| 708766281 | 0 | 16 | 16 | 0 | 0 |
| 716017344 | 0.001759703 | 78 | 0 | 12 | 1 |
| 660448645 | 0.224955155 | 502252 | 463906 | 0 | 0 |
| 667164845 | 0.007239066 | 592 | 97 | 12 | 0 |
| 10935489049 | 1.73746E-05 | 47 | 7 | 21 | 0 |
| 276795119 | 0.041715624 | 20 | 13 | 0 | 0 |
| 1378821457 | 2.650059054 | 4697 | 3860 | 14 | 0 |
| 1260042467 | 0.536421601 | 29676 | 3041 | 0 | 0 |
| 286272949 | 27.18491417 | 99466 | 13755 | 0 | 1 |
| 16461565551 | 0.007467177 | 8319 | 7720 | 19 | 0 |
| 15930099777 | 0.014349382 | 3013 | 2080 | 19 | 0 |
| 6702636467 | 0.000158146 | 348 | 235 | 7 | 0 |
| 471802747 | 1.160084126 | 118125 | 99288 | 0 | 0 |
| 831513739 | 0.002898004 | 302 | 41 | 0 | 0 |
| 812138927 | 0.117870268 | 7736 | 2915 | 8 | 1 |
| 1914824379 | 0.000819912 | 165 | 0 | 7 | 1 |
| 6787317308 | 0.258171028 | 103458 | 9537 | 80 | 0 |
| 736065601 | 12.06045568 | 180720 | 28446 | 15 | 0 |
| 682808182 | 0.009957881 | 675 | 332 | 9 | 1 |
| 552145369 | 1.039059873 | 318 | 203 | 9 | 0 |
| 793463571 | 0 | 28 | 0 | 28 | 0 |
| 303163251 | 0.000857616 | 35 | 0 | 9 | 0 |
| 370221752 | 0.009722956 | 204 | 13 | 11 | 0 |
| 834896214 | 0.005269838 | 887 | 479 | 8 | 2 |
| 2683993746 | 0.125924349 | 35440 | 1909 | 15 | 0 |

| | | | | | |
|---|---|---|---|---|---|
| 373729396 | 0 | 11 | 0 | 11 | 0 |
| 607461963 | 5.3172E-05 | 46 | 13 | 29 | 0 |
| 270080649 | 0.049036843 | 450 | 121 | 14 | 0 |
| 558355857 | 6.111777516 | 76355 | 35436 | 11 | 0 |
| 1194048565 | 0.045816163 | 182 | 78 | 0 | 0 |
| 889060082 | 4.760100677 | 2522 | 1526 | 18 | 0 |
| 968548666 | 4.235418334 | 1204909 | 300377 | 0 | 1 |
| 271732855 | 26.52 | 47483 | 17764 | 9 | 2 |
| 306219677 | 5.70 | 7742 | 430 | 10 | 1 |

| Contig_N50 | Contig_L50 | Scaffold_N50 | Scaffold_L50 | Scaffold_N50/Contig_N50 |
|---|---|---|---|---|
| 1310030 | 78 | 68658214 | 5 | 0.019080456 |
| 419290 | 650 | 20441467 | 18 | 0.020511737 |
| 255416 | 250 | 27368013 | 4 | 0.009332647 |
| 91439 | 2613 | 510878 | 506 | 0.178984024 |
| 69630691 | 15 | 610333535 | 4 | 0.114086294 |
| 29235848 | 27 | 251527947 | 5 | 0.116233001 |
| 139725 | 1033 | 23441323 | 11 | 0.005960628 |
| 12524908 | 55 | 324658466 | 4 | 0.03857872 |
| 10217643 | 364 | 1574410792 | 4 | 0.00648982 |
| 25835044 | 4 | 25835044 | 4 | 1 |
| 26899771 | 9 | 26899771 | 9 | 1 |
| 30060613 | 14 | 53737150 | 10 | 0.559400954 |
| 37717778 | 8 | 37717778 | 8 | 1 |
| 10465369 | 14 | 30457480 | 7 | 0.343605873 |
| 1493114 | 464 | 135027066 | 9 | 0.011057887 |
| 8937712 | 9 | 31125843 | 4 | 0.287147628 |
| 1287207 | 119 | 61987056 | 5 | 0.020765739 |
| 4839000 | 32 | 27625376 | 8 | 0.175165037 |
| 122789 | 6630 | 227195441 | 7 | 0.000540455 |
| 239236 | 1923 | 128314321 | 7 | 0.001864453 |
| 1787824 | 209 | 3844283 | 102 | 0.465060455 |
| 1734059 | 126 | 1734059 | 126 | 1 |
| 19407 | 9578 | 126020 | 1407 | 0.153999365 |
| 78837527 | 5 | 78837527 | 5 | 1 |
| 47037903 | 16 | 226353449 | 5 | 0.207807317 |
| 1885876 | 73 | 1885876 | 73 | 1 |
| 8969346 | 119 | 523395090 | 4 | 0.017136855 |
| 20171672 | 13 | 69564429 | 6 | 0.289971071 |
| 10911736 | 16 | 41684185 | 6 | 0.261771605 |
| 21920 | 5927 | 48366697 | 5 | 0.000453204 |
| 3258867 | 56 | 29151420 | 10 | 0.111791021 |
| 33721 | 4126 | 65803235 | 4 | 0.000512452 |
| 2005037 | 448 | 176490873 | 8 | 0.01136057 |
| 897195 | 196 | 4728343 | 19 | 0.18974829 |
| 6396877 | 139 | 141269392 | 10 | 0.045281408 |
| 1580367 | 177 | 48209797 | 9 | 0.032781034 |
| 23783150 | 12 | 151450279 | 3 | 0.157036026 |
| 31914 | 6191 | 1344915 | 160 | 0.023729381 |
| 32942340 | 4 | 32942340 | 4 | 1 |
| 54239 | 1839 | 36653616 | 5 | 0.001479772 |
| 46553 | 7625 | 131339754 | 5 | 0.000354447 |
| 52169 | 1544 | 2060396 | 44 | 0.02531989 |
| 1072962 | 146 | 37114715 | 7 | 0.028909342 |
| 22561 | 7470 | 557663 | 69 | 0.040456333 |
| 2974952 | 62 | 35530340 | 8 | 0.083729905 |
| 108836 | 618 | 9833969 | 11 | 0.011067352 |
| 48311 | 5797 | 12567911 | 23 | 0.003843996 |

| | | | | |
|---|---|---|---|---|
| 31111469 | 6 | 31111469 | 6 | 1 |
| 341253 | 12216 | 713360525 | 10 | 0.000478374 |
| 1773300 | 436 | 204470928 | 6 | 0.008672627 |
| 3579047 | 35 | 17652500 | 10 | 0.202750149 |
| 23500774 | 12 | 76975026 | 5 | 0.305303879 |
| 67047 | 12505 | 1388941 | 692 | 0.048272029 |
| 28779 | 2191 | 219566 | 316 | 0.131072206 |
| 715870 | 281 | 949204 | 217 | 0.754179291 |
| 114399 | 833 | 11759267 | 13 | 0.009728412 |
| 6076493 | 19 | 51152908 | 4 | 0.118790764 |
| 115182 | 511 | 24375383 | 4 | 0.004725341 |
| 21185935 | 41 | 186523490 | 8 | 0.113583201 |
| 15310187 | 10 | 39611093 | 6 | 0.386512612 |
| 2435 | 45410 | 2435 | 45410 | 1 |
| 400000 | 1486 | 156137564 | 7 | 0.002561843 |
| 13636 | 5014 | 21988864 | 5 | 0.000620132 |
| 2043694 | 48 | 24737479 | 6 | 0.08261529 |
| 28606889 | 8 | 66866305 | 4 | 0.427822189 |
| 15734411 | 48 | 27720101 | 32 | 0.567617376 |
| 3089666 | 29 | 21537881 | 6 | 0.143452645 |
| 32867627 | 7 | 32867627 | 7 | 1 |
| 56484001 | 5 | 56484001 | 5 | 1 |
| 203885 | 461 | 420659 | 240 | 0.48467999 |
| 44734305 | 8 | 44734305 | 8 | 1 |
| 25027151 | 9 | 59921945 | 5 | 0.417662527 |
| 3410 | 49686 | 3565 | 47932 | 0.956521739 |
| 2390495 | 90 | 55380075 | 6 | 0.043165254 |
| 4.3E+08 | 10 | 508132858 | 10 | 0.845294767 |
| 30419810 | 4 | 37865532 | 4 | 0.803364125 |
| 1102861 | 192 | 2981833 | 53 | 0.369860083 |
| 108491 | 3409 | 155806887 | 4 | 0.000696317 |
| 3558 | 24279 | 145010 | 760 | 0.024536239 |
| 94486194 | 57 | 1056413696 | 11 | 0.089440523 |
| 52991057 | 84 | 1056413696 | 11 | 0.050161274 |
| 1.28E+08 | 16 | 943691929 | 4 | 0.135635553 |
| 10988 | 10311 | 18808 | 3416 | 0.584219481 |
| 5393363 | 49 | 35828999 | 11 | 0.150530664 |
| 206950 | 1144 | 94913092 | 4 | 0.002180416 |
| 2.84E+08 | 4 | 283920974 | 4 | 1 |
| 87345 | 22099 | 82862972 | 32 | 0.00105409 |
| 6504 | 34201 | 41463214 | 10 | 0.000156862 |
| 24748136 | 10 | 36467798 | 9 | 0.678629842 |
| 14296461 | 11 | 55506693 | 5 | 0.257562831 |
| 28260955 | 13 | 28260955 | 13 | 1 |
| 16784481 | 7 | 30696496 | 4 | 0.546788174 |
| 3818138 | 30 | 35418074 | 5 | 0.107801966 |
| 3545883 | 72 | 102863307 | 4 | 0.034471797 |
| 127111 | 6056 | 174886800 | 7 | 0.000726819 |

| | | | | |
|---|---|---|---|---|
| 36073259 | 5 | 36073259 | 5 | 1 |
| 20705581 | 13 | 21144679 | 13 | 0.979233641 |
| 10511766 | 10 | 19552154 | 7 | 0.537627005 |
| 21795 | 7711 | 53895704 | 6 | 0.000404392 |
| 18019306 | 28 | 18929811 | 27 | 0.951900999 |
| 1926153 | 101 | 59341207 | 7 | 0.032458945 |
| 1888 | 91315 | 9442 | 26199 | 0.199957636 |
| 10609 | 7109 | 1089885 | 91 | 0.009734055 |
| 87341 | 879 | 36364294 | 5 | 0.002401834 |

| Genome_Coverage | Annotatior | nnotation_Release_Date | Count_Gene_Total | Count_Gene_Protein-codi |
|---|---|---|---|---|
| 8 | 7.4 | 02-06-17 | 32945 | 28236 |
| 8.05 | 10.3 | 28-01-25 | 59048 | 47068 |
| 8.47 | 7.3 | 07-03-17 | 26412 | 23135 |
| 10 | | 02-03-24 | 55405 | 53391 |
| 25 | 9 | 02-11-21 | 58438 | 31448 |
| 32 | 10.2 | 04-01-24 | 57465 | 35582 |
| 34 | 8.5 | 28-01-21 | 35061 | 24562 |
| 35 | 10.1 | 16-12-22 | 51651 | 36855 |
| 38 | | 28-02-25 | 37398 | 37398 |
| 40 | | 08-07-20 | 52342 | 30573 |
| 40 | 10.2 | 14-09-23 | 29591 | 25187 |
| 40 | 10.3 | 27-02-25 | 80883 | 50280 |
| 42 | 10.3 | 19-12-24 | 51804 | 39340 |
| 44 | 10 | 07-10-22 | 28628 | 20741 |
| 48 | 10.3 | 27-02-25 | 110961 | 65994 |
| 50 | 8.3 | 10-12-19 | 23999 | 20040 |
| 50 | 10.1 | 07-06-23 | 30391 | 24186 |
| 51 | 10.3 | 18-09-24 | 42167 | 34369 |
| 56 | 9 | 14-03-22 | 48341 | 32202 |
| 60 | 10.3 | 14-04-25 | 37671 | 28077 |
| 60 | 7.4 | 17-07-17 | 58882 | 49222 |
| 80 | | 10-11-20 | 67843 | 61938 |
| 80 | 8 | 17-05-18 | 30275 | 26313 |
| 82 | | 28-06-25 | 40252 | 40252 |
| 83 | 10.3 | 14-02-25 | 49888 | 34313 |
| 83.15 | 10.3 | 16-09-24 | 34165 | 29679 |
| 85.2 | 10.3 | 07-04-25 | 65673 | 40030 |
| 87 | 10.3 | 20-09-24 | 41476 | 28611 |
| 91 | 8.1 | 11-02-19 | 34292 | 28223 |
| 94 | 6.4 | 20-08-15 | 53125 | 44382 |
| 97 | 9 | 20-09-21 | 34312 | 29735 |
| 97.95 | 10.3 | 18-04-25 | 29938 | 24841 |
| 100 | 8.5 | 11-08-20 | 83308 | 57237 |
| 100 | 8.5 | 19-01-21 | 36764 | 29239 |
| 100 | 9 | 15-07-21 | 89817 | 68154 |
| 100 | 9 | 04-06-22 | 123214 | 90897 |
| 109 | 10.3 | 11-04-25 | 39719 | 28771 |
| 114 | 10.3 | 03-04-25 | 32873 | 28404 |
| 117 | 10 | 05-12-22 | 29331 | 23556 |
| 117 | 10.2 | 14-09-23 | 30909 | 26896 |
| 123 | 7.3 | 24-02-17 | 32237 | 26460 |
| 152.7 | 7.3 | 30-03-17 | 26667 | 24075 |
| 155 | 8.5 | 27-07-20 | 37996 | 30709 |
| 160 | 8.2 | 23-05-19 | 33324 | 29045 |
| 180 | | 14-05-24 | 26098 | 26098 |
| 198 | 8 | 22-01-18 | 35783 | 29279 |
| 220 | 7.4 | 09-11-17 | 47911 | 40126 |

| | | | | |
|---|---|---|---|---|
| 220 | 10.3 | 21-06-24 | 39387 | 29427 |
| 234 | 10.3 | 16-04-25 | 155467 | 103793 |
| 239 | 8.1 | 24-09-18 | 91581 | 63018 |
| 240 | 9 | 15-10-21 | 38372 | 32071 |
| 299 | 10.2 | 16-11-23 | 35194 | 28747 |
| 300 | 8.1 | 26-02-19 | 68531 | 50838 |
| 327 | 7.4 | 25-07-17 | 30405 | 25841 |
| 374 | 8.2 | 18-10-19 | 42351 | 32030 |
| 400 | 7.3 | 13-02-17 | 25758 | 22251 |
| 556 | 10.2 | 08-03-24 | 39086 | 33079 |
| 800 | 8.4 | 13-05-20 | 26936 | 23151 |
| 100 | | | | |
| 100 | | | | |
| 108 | | | | |
| 114 | | | | |
| 116 | | | | |
| 118 | | | | |
| 122.98 | | | | |
| 14 | | | | |
| 147 | | | | |
| 167 | | | | |
| 181.77 | | | | |
| 20 | | | | |
| 20 | | | | |
| 200 | | | | |
| 21 | | | | |
| 243 | | | | |
| 25 | | | | |
| 260 | | | | |
| 288 | | | | |
| 373.57 | | | | |
| 40 | | | | |
| 40 | | | | |
| 40 | | | | |
| 47 | | | | |
| 50 | | | | |
| 53 | | | | |
| 54 | | | | |
| 60 | | | | |
| 66 | | | | |
| 67 | | | | |
| 72 | | | | |
| 73.49 | | | | |
| 80 | | | | |
| 82 | | | | |
| 84 | | | | |
| 84 | | | | |
| 85 | | | | |

| | | | | |
|---|---|---|---|---|
| 85 | | | | |
| 86.74 | | | | |
| 899.8 | | | | |
| 90.97 | | | | |
| 95 | | | | |
| 98 | | | | |
| NA | | | | |
| | 7.4 | 03-08-17 | 20332 | 18126 |
| | 7.1 | 05-09-16 | 24957 | 21518 |

| Count_Gene_Non-coding | Count_Gene_Pse | Annotation_BUSCO_Lineage |
|---|---|---|
| 3461 | 1248 | poales_odb10 |
| 7219 | 4761 | fabales_odb10 |
| 1893 | 1384 | eudicots_odb10 |
| 1210 | 80 | |
| 21210 | 5778 | poales_odb10 |
| 19381 | 2502 | eudicots_odb10 |
| 7923 | 2576 | liliopsida_odb10 |
| 12130 | 2666 | eudicots_odb10 |
| 2953 | 1451 | eudicots_odb10 |
| 25576 | 5027 | eudicots_odb10 |
| 9627 | 2837 | eudicots_odb10 |
| 7357 | 530 | eudicots_odb10 |
| 40614 | 4353 | fabales_odb10 |
| 3589 | 370 | eudicots_odb10 |
| 5056 | 1035 | eudicots_odb10 |
| 3725 | 4073 | eudicots_odb10 |
| 10390 | 5748 | solanales_odb10 |
| 5557 | 4037 | liliopsida_odb10 |
| 6105 | 3555 | eudicots_odb10 |
| | 5905 | |
| 2877 | 1085 | |
| 10353 | 5222 | poales_odb10 |
| 3074 | 1412 | fabales_odb10 |
| 20260 | 5383 | fabales_odb10 |
| 11404 | 1457 | solanales_odb10 |
| 4855 | 1214 | |
| 4248 | 4495 | brassicales_odb10 |
| 3366 | 1208 | eudicots_odb10 |
| 3736 | 1361 | fabales_odb10 |
| 23778 | 2293 | eudicots_odb10 |
| 5389 | 2136 | liliopsida_odb10 |
| 16202 | 5461 | liliopsida_odb10 |
| 23012 | 9304 | brassicales_odb10 |
| 9543 | 1405 | eudicots_odb10 |
| 2434 | 2027 | solanales_odb10 |
| 4524 | 1251 | eudicots_odb10 |
| 2042 | 1971 | eudicots_odb10 |
| 4229 | 1548 | liliopsida_odb10 |
| 1998 | 594 | eudicots_odb10 |
| 4930 | 2357 | eudicots_odb10 |
| 3093 | 1186 | |
| 5719 | 785 | |
| 4367 | 3418 | eudicots_odb10 |

| | | |
|---|---|---|
| 8050 | 1910 | poales_odb10 |
| 32609 | 19065 | poales_odb10 |
| 19945 | 8618 | embryophyta_odb10 |
| 3683 | 2618 | eudicots_odb10 |
| 5506 | 941 | eudicots_odb10 |
| 11498 | 6195 | eudicots_odb10 |
| 2959 | 1605 | eudicots_odb10 |
| 4632 | 5689 | eudicots_odb10 |
| 2930 | 577 | poales_odb10 |
| 5090 | 917 | eudicots_odb10 |
| 2292 | 1493 | eudicots_odb10 |

| | | |
|---|---|---|
| 1874 | 332 | brassicales_odb10 |
| 2274 | 1165 | eudicots_odb10 |

| BUSCO_Complete | BUSCO_Single_Copy | BUSCO_Duplicated | BUSCO_Fragmented |
|---|---|---|---|
| 0.9965278 | 0.98386437 | 0.012663399 | 0.000612745 |
| 0.99180025 | 0.3911666 | 0.6006336 | 0.003168095 |
| 0.99312127 | 0.97936374 | 0.013757524 | 0.001719691 |
| 0.98672384 | 0.95547384 | 0.03125 | 0.002655229 |
| 0.9853826 | 0.9522786 | 0.03310404 | 0.000859845 |
| 0.9242892 | 0.8615575 | 0.062731765 | 0.050061803 |
| 0.9840929 | 0.9505589 | 0.033533964 | 0.002149613 |
| 0.9879622 | 0.97248495 | 0.015477214 | 0.002149613 |
| 0.9840929 | 0.059759244 | 0.92433363 | 0.005588994 |
| 0.9849527 | 0.5950129 | 0.3899398 | 0.004729149 |
| 0.98194325 | 0.96861565 | 0.013327601 | 0.003009458 |
| 0.9754007 | 0.05273947 | 0.9226612 | 0.003354454 |
| 0.9810834 | 0.9531384 | 0.027944969 | 0.003439381 |
| 0.96689594 | 0.94411004 | 0.022785898 | 0.003439381 |
| 0.9789338 | 0.61306965 | 0.36586416 | 0.005159071 |
| 0.9890756 | 0.9628571 | 0.026218487 | 0.002857143 |
| 0.9449938 | 0.7839926 | 0.16100124 | 0.034301605 |
| 0.98366296 | 0.1177988 | 0.86586416 | 0.001719691 |
| 0.9781454 | 0.8251634 | 0.15298203 | 0.005514706 |
| 0.98360044 | 0.96515095 | 0.018449496 | 0.00260902 |
| 0.9793142 | 0.93458813 | 0.04472605 | 0.004099888 |
| 0.98689073 | 0.9680672 | 0.018823529 | 0.001344538 |
| 0.9945605 | 0.8996954 | 0.0948651 | 0.001958225 |
| 0.99527085 | 0.9183147 | 0.076956145 | 0.002149613 |
| 0.9806187 | 0.96477824 | 0.015840476 | 0.004472605 |
| 0.9759243 | 0.8417885 | 0.13413586 | 0.002149613 |
| 0.9576638 | 0.7673053 | 0.19035846 | 0.02750309 |
| 0.9527194 | 0.09023486 | 0.8624846 | 0.03275649 |
| 0.99695385 | 0.042210616 | 0.95474327 | 0.000435161 |
| 0.9733448 | 0.938951 | 0.03439381 | 0.005159071 |
| 0.9882353 | 0.96789914 | 0.020336134 | 0.00302521 |
| 0.9922614 | 0.9690456 | 0.023215821 | 0.002149613 |
| 0.9892519 | 0.92304385 | 0.06620808 | 0.004299226 |
| 0.8800989 | 0.8399258 | 0.040173054 | 0.09332509 |
| 0.9849527 | 0.9282029 | 0.056749783 | 0.003439381 |
| 0.99269134 | 0.9011178 | 0.091573514 | 0.001719691 |
| 0.85855544 | 0.72398967 | 0.13456579 | 0.058469474 |

| | | | |
|---|---|---|---|
| 0.9930556 | 0.9738562 | 0.019199347 | 0.002246732 |
| 0.99611926 | 0.010825164 | 0.9852941 | 0.001429739 |
| 0.988228 | 0.1685254 | 0.8197026 | 0.002478315 |
| 0.9961307 | 0.7901978 | 0.20593293 | 0.001289768 |
| 0.9823732 | 0.92132413 | 0.06104901 | 0.000859845 |
| 0.9638865 | 0.83190024 | 0.13198625 | 0.011607911 |
| 0.9797936 | 0.9466896 | 0.03310404 | 0.007738607 |
| 0.96474636 | 0.7742906 | 0.19045572 | 0.015477214 |
| 0.93852127 | 0.87009805 | 0.068423204 | 0.007965687 |
| 0.99140155 | 0.94196045 | 0.0494411 | 0.000859845 |
| 0.977644 | 0.92175406 | 0.055889938 | 0.002579536 |

| | | | |
|---|---|---|---|
| 0.8083116 | 0.7822019 | 0.02610966 | 0.097911224 |
| 0.99785036 | 0.9909716 | 0.006878762 | 0.000859845 |

| BUSCO_Missing | Genome LAI |
|---|---|
| 0.002859477 | |
| 0.005031681 | |
| 0.005159071 | |
| 0.010620915 | 12.9 |
| 0.013757524 | 11.1 |
| 0.02564895 | |
| 0.013757524 | 19.66 |
| 0.009888221 | 12.8 |
| 0.010318142 | |
| 0.010318142 | 13.6 |
| 0.015047291 | 11.2 |
| 0.021244874 | |
| 0.015477214 | |
| 0.02966466 | |
| 0.015907137 | 10.9 |
| 0.008067227 | |
| 0.020704573 | |
| 0.014617369 | |
| 0.01633987 | 13.3 |
| 0.013790533 | 18.61 |
| 0.01658591 | 17.5 |
| 0.011764706 | 12.7 |
| 0.003481288 | |
| 0.002579536 | 18.79 |
| 0.014908684 | |
| 0.021926053 | |
| 0.014833127 | |
| 0.014524104 | |
| 0.002610966 | |
| 0.02149613 | 20 |
| 0.008739496 | |
| 0.005588994 | 20.68 |
| 0.006448839 | 17.7 |
| 0.02657602 | |
| 0.011607911 | |
| 0.005588994 | |
| 0.08297507 | |

| | |
|---|---|
| 0.004697713 | 20.3 |
| 0.002450981 | |
| 0.00929368 | |
| 0.002579536 | 12.01 |
| 0.016766982 | |
| 0.02450559 | |
| 0.012467756 | |
| 0.019776441 | |
| 0.053513072 | |
| 0.007738607 | 15.03 |
| 0.019776441 | |

| |
|---|
| 0.093777195 |
| 0.001289768 |
