## Supplementary File 1 for "GAP-MS: Automated validation of gene predictions using integrated mass spectrometry evidence"

### Supplementary File 1: GAP-MS: Automated validation of gene predictions using integrated mass spectrometry evidence

#### XGBoost Model training and performance evaluation

An XGBoost (eXtreme Gradient Boosting) classifier was developed to classify gene models with intermediate proteomics evidence as either verified or dismissed. The model captures nonlinear relationships between multiple proteomics-derived features that would be systematically misclassified using rule-based filtering approaches alone.

##### 1. Training data preparation

The training and validation dataset consisted of two classes:

- **Positive examples**: Gene models from the high-confidence set, defined as features satisfying all rule-based filtering criteria indicative of genuine gene models. This set represented well-supported gene models with strong proteomics evidence.
- **Negative examples:** Gene models from the low-confidence set, defined as features failing one or more rule-based criteria, representing likely erroneous predictions or artifacts.
- **Test set (Unlabeled)**: Gene models with intermediate proteomics evidence that did not meet all high-confidence criteria nor fully satisfy low-confidence exclusion criteria. This unlabeled set required probabilistic classification.

##### 2. Feature engineering and model input variables

The XGBoost classifier utilized four primary proteomics-derived features:

- **Number of mapped peptides** to the protein sequence.
- **Sequence coverage** is defined as the percentage of the protein sequence validated by peptide matches.
- **Peptide uniqueness** is the proportion of peptides uniquely mapping to this gene model vs. mapping to multiple loci.
- **Peptide location**: the positional diversity of peptide evidence across the protein sequence; measured as coefficient of variation of peptide start positions.

All features were standardized using z-score normalization (zero mean, unit variance) prior to model training. Pairwise feature correlations were examined to identify potential multicollinearity. Feature importance rankings were derived from the trained model to assess the relative contribution of each variable to classification decisions.

##### 3. Model architecture and hyperparameters

An XGBoost classifier was used for binary classification with probabilistic outputs (binary logistic loss). The base learners were shallow decision trees.

Hyperparameters were optimized using randomized search over the following ranges:

| | | |
|---|---|---|
| n_estimators | 50-500 | Number of boosting rounds/trees |
| max_depth | 3-15 | Maximum tree depth |
| learning_rate | 0.001-0.3 | Shrinkage parameter controlling step size |

| | | |
|---|---|---|
| min_child_weight | 1-10 | Minimum sum of instance weights in child nodes |
| subsample | 0.5-1.0 | Fraction of training samples used per iteration |
| colsample_bytree | 0.5-1.0 | Fraction of features sampled per tree |
| gamma | 0.0-5.0 | Minimum loss reduction for split |
| lambda | 0-2.0 | L2 regularization parameter |
| alpha | 0-2.0 | L1 regularization parameter |

**4. Cross-validation, Hyperparameter search, and early stopping**

Model evaluation used stratified 5-fold cross-validation to preserve class distribution across folds and ensure robust performance estimation.

For each fold, 80% of the data was used for training and 20% for validation. Reported performance metrics represent the mean ± standard deviation across the five folds.

Randomized hyperparameter search sampled 100 parameter combinations from the defined search space and evaluated them using stratified 5-fold cross-validation. The configuration with the highest mean ROC-AUC was selected.

To reduce overfitting and training time, early stopping terminated training if validation performance did not improve for 10 consecutive rounds.

**5. Performance evaluation metrics**

Model performance was assessed using multiple complementary metrics appropriate for binary classification with imbalanced datasets:

**Primary metric**: Receiver Operating Characteristic - Area Under Curve (ROC-AUC) was selected as the primary optimization objective for hyperparameter search.

Performance was further characterized using:

- Accuracy: (TP + TN) / (TP + TN + FP + FN)
- Sensitivity (Recall): TP / (TP + FN)
- Specificity: TN / (TN + FP)
- Precision: TP / (TP + FP)
- F1-Score

*Where TP = true positives, TN = true negatives, FP = false positives, FN = false negatives.*

For detailed assessment of classification performance, confusion matrices were constructed for each cross-validation fold to evaluate the distribution of correct and incorrect classifications.

**8. Final model application**

The final optimized XGBoost model, retrained on the complete training dataset (high-confidence and low-confidence sets combined), was applied to the unlabeled set of gene models with intermediate evidence. The model produced probability scores for each gene model, representing the predicted likelihood of being verified.
